# Allosteric regulation of lysosomal enzyme recognition by the cation-independent mannose 6-phosphate receptor

**DOI:** 10.1101/2020.02.21.959957

**Authors:** Linda J. Olson, Sandeep K. Misra, Mayumi Ishihara, Kevin P. Battaile, Oliver C. Grant, Amika Sood, Robert J. Woods, Jung-Ja P. Kim, Michael Tiemeyer, Gang Ren, Joshua S. Sharp, Nancy M. Dahms

**Affiliations:** Department of Biochemistry, Medical College of Wisconsin, Milwaukee, WI; Department of BioMolecular Sciences, University of Mississippi, Oxford, MS; The Molecular Foundry, Lawrence Berkeley National Laboratory, Berkeley, CA; Complex Carbohydrate Research Center, University of Georgia, Athens, GA; IMCA-CAT, Hauptman-Woodward Medical Research Institute, Argonne, IL; current affiliation New York Structural Biology Center, New York City, NY

## Abstract

The cation-independent mannose 6-phosphate receptor (CI-MPR), also known as the IGF2 receptor or CD222, is a multifunctional type I transmembrane glycoprotein ubiquitously expressed in most eukaryotic cell types. Through the receptor’s ability to bind a variety of unrelated extracellular and intracellular ligands, it is involved in a wide array of functions including protein trafficking, lysosomal biogenesis, internalization, regulation of cell growth, cell migration and apoptosis. CI-MPR has a large extracellular region comprised of 15 contiguous domains, four of which interact with phosphorylated glycans on lysosomal enzymes. Here we present a series of biophysical studies, along with crystal structures, providing information on how the N-terminal 5 domains of this receptor work in concert to bind and release carbohydrates. High-resolution electron microscopy as well as hydroxyl radical protein footprinting (HRPF) of this multifunctional multidomain construct demonstrates dynamic conformational changes occur as a consequence of ligand binding and different pH conditions, These data, coupled with surface plasmon resonance studies and molecular modeling, allow us to propose a bi-dentate oligosaccharide binding model, which could explain how high affinity carbohydrate binding is achieved through allosteric domain cooperativity.

## Introduction

Lysosomes are acidified organelles that carry out degradative metabolism critical to many endocytic, phagocytic, and autophagic processes, such as inactivation of pathogenic organisms and disposal of abnormal proteins^1–5^. This diverse degradative capacity depends on a collection of over 60 different soluble proteases, glycosidases, nucleases, and lipases. Delivery of these newly synthesized hydrolytic enzymes to lysosomes depends on the P-type lectins, the 300kDa cation-independent mannose 6-phosphate receptor (CI-MPR) and the 46kDa cation-dependent MPR (CD-MPR), that bind a unique carbohydrate determinant, mannose 6-phosphate (M6P), on lysosomal enzymes. However, CI-MPR is the primary receptor responsible for this trafficking^4^. Because CI-MPR binds a wide range of ligands^6,7^ at the cell surface that include M6P-containing cytokines/hormones^8,9^ and non-M6P-containing molecules (*e.g*., insulin-like growth factor 2 (IGF2)^10^, plasminogen^11^, urokinase-type plasminogen activator receptor (uPAR)^11^) to mediate CI-MPR’s roles as a tumor suppressor^12^ and regulator of cell growth and differentiation^13^, it is not surprising CI-MPR is essentiel for normal development as transgenic mice lacking the CI-MPR gene die at birth^14,15^.

Lysosomal storage diseases (LSDs) are caused by mutations in lysosomal proteins, mainly enzymes, that result in defective catabolism and substrate accumulation. Characteristic of the family of ~70 LSDs is their progressive and debilitating nature due to their impact on multiple organ systems. Treatment is symptomatic for most LSDs, with only 11 having FDA-approved therapies. For example, deficiency of palmitoyl-protein thioesterase 1 (PPT1), which removes thioester-linked fatty acyl groups from cysteine residues of lipid-modified proteins, causes the fatal neurodegenerative disorder infantile neuronal ceroid lipofuscinosis. This LSD is characterized by cognitive and motor deterioration that leads to a chronic vegetative state and early death, and currently there are no FDA-approved treatments for these infants^16^. CI-MPR’s ability to internalize recombinant M6P-containing enzymes delivered to patients by bi-weekly intravenous infusion forms the basis of enzyme replacement therapy (ERT) for 9 of these FDA-approved therapies^17–19^. Despite CI-MPR’s critical function in supplying lysosomes with hydrolases and its role in human therapies, knowledge of how CI-MPR interacts with a heterogeneous population of ~60 different lysosomal enzymes is lacking, and no structure of CI-MPR, or CD-MPR, bound to an enzyme is currently available.

Many lectins bind sugars by simultaneously engaging multiple sugar binding sites, termed carbohydrate recognition domains (CRDs), that can be located on a single polypeptide chain (tandem repeats of CRDs) or on different polypeptide chains (hetero-oligomers or clustering of monomers on the cell surface). The resulting multivalent interactions serve to greatly increase ligand affinity^20^. CI-MPR’s large extracytoplasmic region contains 15 homologous domains called ‘MRH’ domains (<u>M</u>annose 6-phosphate <u>R</u>eceptor <u>H</u>omology) due to their similar size (~150 residues) and conserved residues, including cysteines involved in disulfide bonding^21^. The binding sites for its diverse ligands map to different MRH domains of this multifunctional receptor. For example, IGF2 binds to domain 11 via protein-protein interactions^22^. Additionally, we showed CI-MPR has four non-adjacent CRDs (domains 3, 5, 9, and 15), each with distinctive phosphomonoester and phosphodiester glycan preferences leading to high affinity binding (Fig. 1a)^23–25^. Our crystal and NMR structures of several MRH domains of CI-MPR (domains 1-3, domain 5)^26,27^, along with the structures solved by Jones *et al.* of CI-MPR’s domains 11-14^28–30^, reveal that each MRH domain has a similar β-barrel fold comprised of two β-sheets positioned orthogonally over each other. However, a predictive structural model of CI-MPR’s entire extracellular region is not possible because in the known multi-domain structures, the assembly of domains differs: the N-terminal three domains exhibit a wedged-shaped arrangement^31^ while domains 11-14 adopt an elongated structure^28^ (Fig. 1a). Furthermore, although domain-domain interactions have been shown to stabilize the binding site for IGF2 (domain 11)^22^ and M6P (domain 3)^26^, how these interactions influence the function and overall structural dynamics of CI-MPR is not fully understood.

**Fig. 1.**
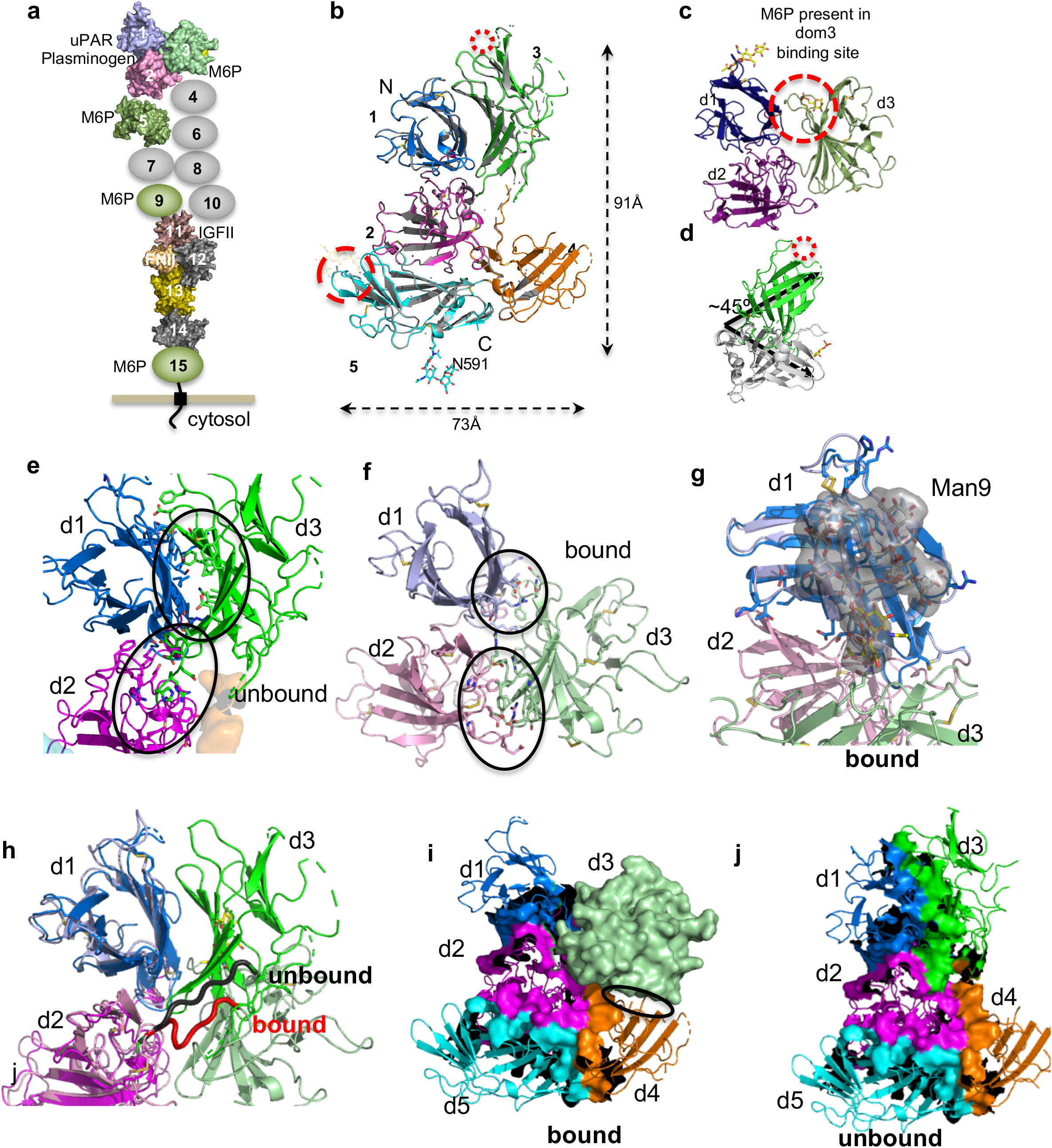
Crystal structure of domains 1-5 of CI-MPR. **a,** Cartoon of domain structure of CI-MPR highlighting multifunctionality of the protein. Relevant ligands are listed next to the known domain of interaction. **b,** Comparison of crystal structures solved at pH 5.5 (PDB 6P8I) and 7.0 (grey)(PDB 6V02). MRH domains displaying the classic fold of two β-sheets orthogonally oriented relative to each other are labeled 1-5 along with the N- and C-terminus. The red star marks the unoccupied known M6P binding site in domain 3, while the glycan of a crystallographic neighbor occupies the known binding site of domain 5 which is circled in red. Approximate dimensions of the domains 1-5 model are shown. **c,** Crystal structure of ligand bound domain (PDB 1SYO) with M6P in the domain 3 binding site (circled). **d,** Comparison of domain 3 in absence and presence of M6P. The empty binding site is circled in the unbound form. **e, f,** Domain structural differences between bound and unbound. Domains are labeled and regions of interest are circled. **g.** Molecular surface of N365 glycosylated with a Man_9_ glycan showing the potential area of protection. **h**, Superimposition of domains 1 of PDB 1SYO (bound domain 3) and PDB 6P8I (unbound domain 3) highlighting the change in linker between the two forms (red, ligand bound domain 3; black, ligand unbound domain 3). **i,** Surface representation of residues forming domain interfaces of the ligand bound orientation of domain 3 (PDB 1SYO) and positions of domains 1-2, and domains 4-5 from PDB 6P8I model**. j,** Surface representation of residues forming domain interfaces of the unbound domain 3 model, PDB 6P8I.

We now report the crystal structure of the N-terminal five domains of human CI-MPR, revealing for the first time the orientation of two CRDs (domain 3 and domain 5) with respect to each other. To our knowledge, this is the first structure of any region of CI-MPR encompassing multiple CRDs, in addition to uPAR and plasminogen binding sites. Analyses of the receptor bound to PPT1 by small angle X-ray scattering (SAXS) and hydroxyl radical protein footprinting (HRPF) and under different pH conditions (e.g., Golgi, late endosome) support binding-induced and pH-induced conformational change. Additionally, high-resolution negative-staining electron microscopy (NS EM) images indicate CI-MPR adopts multiple conformations that are influenced by M6P binding. Furthermore, quantitative binding measurements coupled with biophysical analyses support allosteric regulation of the two CRDs.

### Effects of ligand binding on domain orientation

Crystallization screening of human CI-MPR domains 1-5 protein in the presence of 10 mM M6P resulted in two conditions yielding diffraction quality crystals. Comparison of the two structures, one obtained at ~pH 5.5 (2.5Å, PDB 6P8I) and the other at pH ~7.0 (2.8Å, PDB 6V02), reveals the same domain orientations relative to one another, an inverted ‘T’, with an *r.m.s.d*. of ~0.2 Å over 524 Cα atoms (Fig. 1b, Extended Data Table 1). Importantly, the pH 5.5 condition resulted in a crystal structure with the *N*-glycan at N591 of a crystallographic neighbor occupying the carbohydrate binding site of domain 5, while the carbohydrate binding site in domain 3 is unoccupied. Because all of our previous structures of bovine CI-MPR domains 1-3 show domain 3 bound to ligand, either M6P or the non-phosphorylated mannose oligosaccharide of a crystallographic neighbor^31,32^, the current structure of domains 1-5 reported here allows us the opportunity to evaluate the consequence of carbohydrate binding to a single CRD on individual domain structures as well as the overall positioning of domains relative to one another. Due to the higher resolution and greater degree of completeness, we focus our analyses on the pH ~5.5 structure (PDB 6P8I).

A comparison of the individual domains of our previously published CI-MPR bovine domains 1-3 (PDB 1SYO, ligand-bound domain 3) to the corresponding domains in our current structure of human domains 1-5 (PDB 6P8I, ligand-free domain 3, ligand-bound domain 5) demonstrate that the corresponding individual domains retain the overall core structure (*r.m.s.d*. <0.5 Å). However, there are a few noteworthy differences comparing the bound to unbound structures. First, loop C (between strands 6 and 7) of domain 1 relocates 6.4 Å towards loop C of domain 3 (Extended Data Fig. 1a). Second, the loop between strands 5 and 6 of domain 1 moves towards domain 3 by 4.2 Å allowing it to form contacts with residues in domain 3 (Extended Data Fig. 1a). Third, domain 2 has multiple loops experiencing conformational changes as well as modest changes in the β-strands of the N-terminal (‘front’) sheet relative to the strands of the C-terminal (‘back’) sheet (Extended Data Fig. 1b). The position of domain 2 changes with respect to domain 1 by 0.5 to 2.1 Å (at the junction point between domains 1, 2 and 3) (Extended Data Fig. 1c). Fourth, the Cα of S386, located at the tip of loop C and essential for ligand binding^33^, moves away from the binding cavity by 2.7 Å in the unbound structure (Extended Data Fig 1d). Finally, a dramatic change is observed in the position of domain 3, with an ~45° rotation accompanied by a 34 Å movement of S386 (located on tip of loop C) towards domain 4 (Fig 1c-d). Together, these findings indicate that the N-terminal region of CI-MPR is dynamic, with individual domains altering positions relative to one another depending on the occupancy of domain 3’s carbohydrate binding site.

### Comparison with P-type lectin family member CD-MPR

CI-MPR and CD-MPR are the only members of the P-type lectin family^33^. CD-MPR is a homodimer with a single MRH domain per polypeptide. CD-MPR transitions, through what was previously described as a scissoring motion, from a more closed (binding sites closer together), smaller dimer interface to a more open (larger distance between binding sites) conformation in the presence of M6P (Extended Data Fig. 1e)^34^. These movements triggered by ligand binding increase the size of the interface by ~36% and add two salt bridges, but create an energetically less favorable association (Complex Formation Significance Score (CSS) 1.0 unbound and 0.46 bound as calculated by PISA^35^) (Extended Data Fig. 1f-h). In contrast, domain 3 of CI-MPR has a composite interface made up of interactions with two other domains (domains 1 and 2). Rearrangement of domain 3 upon binding ligand reorients its C-terminal β-sheet to interact with both domains 1 and 2: salt bridge contacts are altered and the interface with domain 1 is reduced by almost two thirds exposing nonpolar residues (Fig. 1e-f, Extended Data Fig. 1f). However, these nonpolar residues may be sheltered from solvent by the presence of a *N*-glycan on domain 3 at position N365, which is species conserved (Fig. 1g, Extended Data Fig. 2)^36^. Given the significant repositioning of domain 3 upon ligand binding, it is not surprising that the linker region connecting domains 2 and 3 changes conformation, adopting a more extended structure in the absence of ligand (Fig. 1h). The ability to alter conformations of this 9-residue linker region appears a key factor in the repositioning of domain 3, and its importance is further reflected in the high species conservation of amino acids of this region (Extended Data Fig. 2). Together, these findings show that the presence of ligand alters the very nature of the relationship of domain 3 to its neighboring domains.

Despite CI-MPR being a multidomain protein and utilizing changes in linker structure to facilitate domain reorientation upon ligand binding, the resulting domain-domain interactions have similarities to its homodimeric family member, CD-MPR: both receptors use residues located on flexible loops to generate salt bridges with the neighboring domain(s) and both use hydrophobic cores comprised of residues within the C-terminal β-sheet. However, CD-MPR maintains a back-to-back (C-terminal β-sheet of one monomer against the C-terminal β-sheet of the other monomer) arrangement in both ligand bound and unbound states (Extended Data Fig. 1e, h), whereas CI-MPR’s domain 1 to domain 3 back-to-back arrangement is found only in the ligand-unbound state (Fig. 1h-j).

### Proposed model of CI-MPR domains 1-5 when both carbohydrate binding sites are occupied

We generated a new model of domains 1-5 by substituting domains 1-3 of PDB 6P8I with the PDB 1SYO structure. Superimposition of domain 1 from these two structures (PDB 6P8I, domains 1-5 with only domain 5 bound, and PDB 1SYO, domains 1-3 with domain 3 bound) shows that the relocation of domain 3 does not introduce major steric clashes with domains 4 and 5 (Fig. 1i), providing us with a plausible view of what the receptor would look like if each of the two carbohydrate binding sites (domain 3 and 5) were simultaneously occupied with ligand. The location of domain 3 in this new model not only serves to significantly increase the interdomain interface area between domains 3 and 4, but the position of domain 4 appears to limit the possible range of motion of domain 3 upon ligand binding (Fig. 1i-j). We carried out additional studies described below to evaluate the validity of this new, proposed model of CI-MPR in which both domains 3 and 5 are bound to ligand.

### Conformation of CI-MPR bound to a lysosomal enzyme

Because we were unsuccessful in obtaining crystals of CI-MPR domains 1-5 with an alternate ligand binding scenario, we turned to small angle X-ray scattering (SAXS) to gather information on effects of 1) ligand binding to domain 3 and 2) ligand absence in domain 5, on the overall structure. Although SAXS provides lower resolution data and represents an overall average of structures in solution, it has proven to be a robust method to explore biomolecular shapes as well as conformational changes under physiological conditions^37^. We collected SAXS data at pH 6.5 in the absence and presence of M6P or PPT1, both which interact preferentially with the phosphomonoester-specific binding site of domain 3^24,38^. Kratky Plots were used to evaluate the protein’s flexibility as well as its globular nature (Extended Data Fig. 3a). Domains 1-5 protein produced plots that are relatively hyperbolic, characteristic of globular proteins, but do have some tailing indicative of the presence of protein flexibility. Additionally, the curves did not significantly change in the presence of either ligand (Extended Data Fig. 3a). Application of Porod-Debye criteria to further assess flexibility show CI-MPR domains 1-5 protein, whether in the presence or absence of ligand, produces a Porod Exponent (P_E_) of 2.8-2.9 that is indicative of a flexible protein with perhaps some intrinsic disorder (Extended Data Fig. 3b,c)^37^.

We then calculated three-dimensional (3D) *ab initio* models of domains 1-5 in the absence and presence of ligand and compared them to our crystallographic models. The overall shape of the calculated envelopes in the presence and absence of the monosaccharide M6P are similar (Extended Data 3d, Fig. 2a) and resembles a sock with a dimple near the heel. Because the small carbohydrate M6P cannot itself be identified by SAXS, we focused our efforts on the human lysosomal enzyme PPT1. This recombinant, monomeric 279-residue protein harbors three *N*-linked glycans and the crystal structure of a monomeric form has already been determined^39^. PPT1 alone gives rise to an oblong 3D model and its position is easily discernable in complex with domains 1-5 (Fig. 2b,c). Based on its preference for the phosphomonoester binding site of domain 3, the envelope is consistent with PPT1 binding to CI-MPR’s domain 3 through its M6P-containing glycans. The 1:1 stoichiometry of the complex was confirmed by calculating the molecular weight from the envelope volume (MW, (in kDa)=V_p_ (in nm^3^)/1.6 ^40^)(Extended Data Fig. 3c). Inspection of the 3D envelopes of domains 1-5 bound either to PPT1 (Fig. 2c) or M6P (Extended Data Fig. 3d) shows lack of molecular model in the toe region, consistent with domain 5, and perhaps domain 4, residing in a different location than that found in our crystallographic structure (PDB 6P8I). Supporting this notion is the observation that domain 4 in our crystallographic structures appears flexible as demonstrated by discontinuous density especially in the pH 7.0 structure (PDB 6V02).

**Fig. 2.**
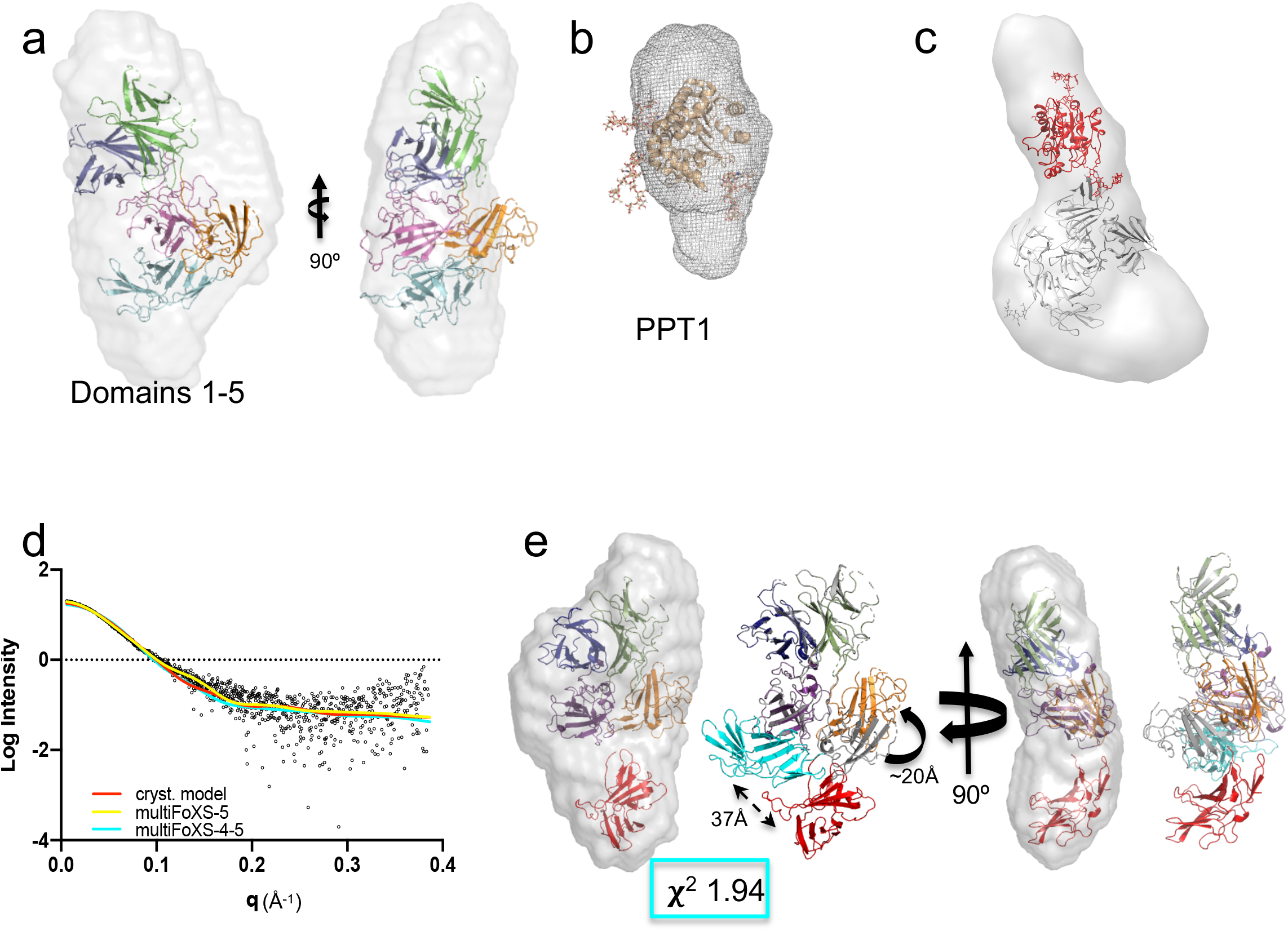
*Ab initio* envelope models rendered as volumes and superimposed onto X-ray crystallographic models. **a**, X-ray model (PDB 6P8I) placed within envelope derived from SEC-SAXS data of domains 1-5 collected in the absence of PPT1 (PDB 1EI9) (**b**). **c**, The modified X-ray model of domains 1-5 (grey), representing domain 5 in the bound position with the PPT1 model (red) (PDB 1EI9) placed within the calculated *ab initio* envelope (rendered as a volume illustrating extra density along the most elongated axis). **d**, Experimental scattering curve for domains 1-5 (black) overlaid with calculated scattering curves generated from X-ray model (PDB 6P8I, red) or mulitFoXS generated model based on PDB 6P8I where either the linker before domain 5 (yellow) or before both domains 4 and 5 (cyan) are allowed to be flexible. Corresponding χ^2^ values for each curve are shown in figures. **e**, MultiFoXS derived model of domains 1-5 in the absence of ligand placed in the same envelope as in (**a)** showing relative movements of domains 4 (grey, PDB 6P8I, to orange) and 5 (cyan, PDB 6P8I, to red).

We next used the program MultiFoXS to model possible orientations of domains 5 and/or 4 to improve the fit of our model to the SAXS scattering curves (Fig. 2d)^41^. Starting with the simplest scenario, only allowing flexibility of the linker between domains 4 and 5, an improved model was calculated with domain 5 swinging into the unpopulated ‘toe’ region (~60 Å) (Extended Data Fig. 3e). Next we allowed flexibility between both domains 3 and 4 as well as domains 4 and 5 (Fig. 2e). In this model, domain 5 has translated into the toe of the SAXS envelope with N682 in loop C of domain 5 translating 37 Å and domain 4 has been displaced from its original position in the crystal structure. Allowing either domain 5 or domains 4 and 5 to be flexible and assume alternate conformations from our crystal structure improved the χ^2^ of the model fit to the scattering curve from 6.62 to 1.47/1.97(Fig. 2d). Together, these SAXS data (Fig. 2d-e, Extended Data Fig. 3e) are consistent with our hypothesis that domains 4 and 5 are flexible and exist in alternate conformations in the absence of ligand in domain 5.

To determine the extent of motions these domains undergo, we negatively stained domains 1-5 in the presence and absence of M6P at pH 7.4 and imaged the samples by high-resolution electron microscopy (EM). The reference-free 2D classifications yielded 50 classes of domain arrangements in the absence of M6P (Fig. 3). The diversity in the negatively-stained EM images demonstrates how dynamic this region of CI-MPR is in the absence of ligand. Although the binding of M6P to domain 3 reduces the number of classes, indicating a reduction in domain mobility upon binding, there are still numerous classes representing multiple conformations of domains 1-5. Together, these SAXS and EM data indicate a high degree of flexibility of CI-MPR’s N-terminal five domains. However, because domain 5 is tethered to the C-terminal domains 6-15 in the native structure, its mobility may be constrained in the context of the full-length receptor.

**Fig. 3.**
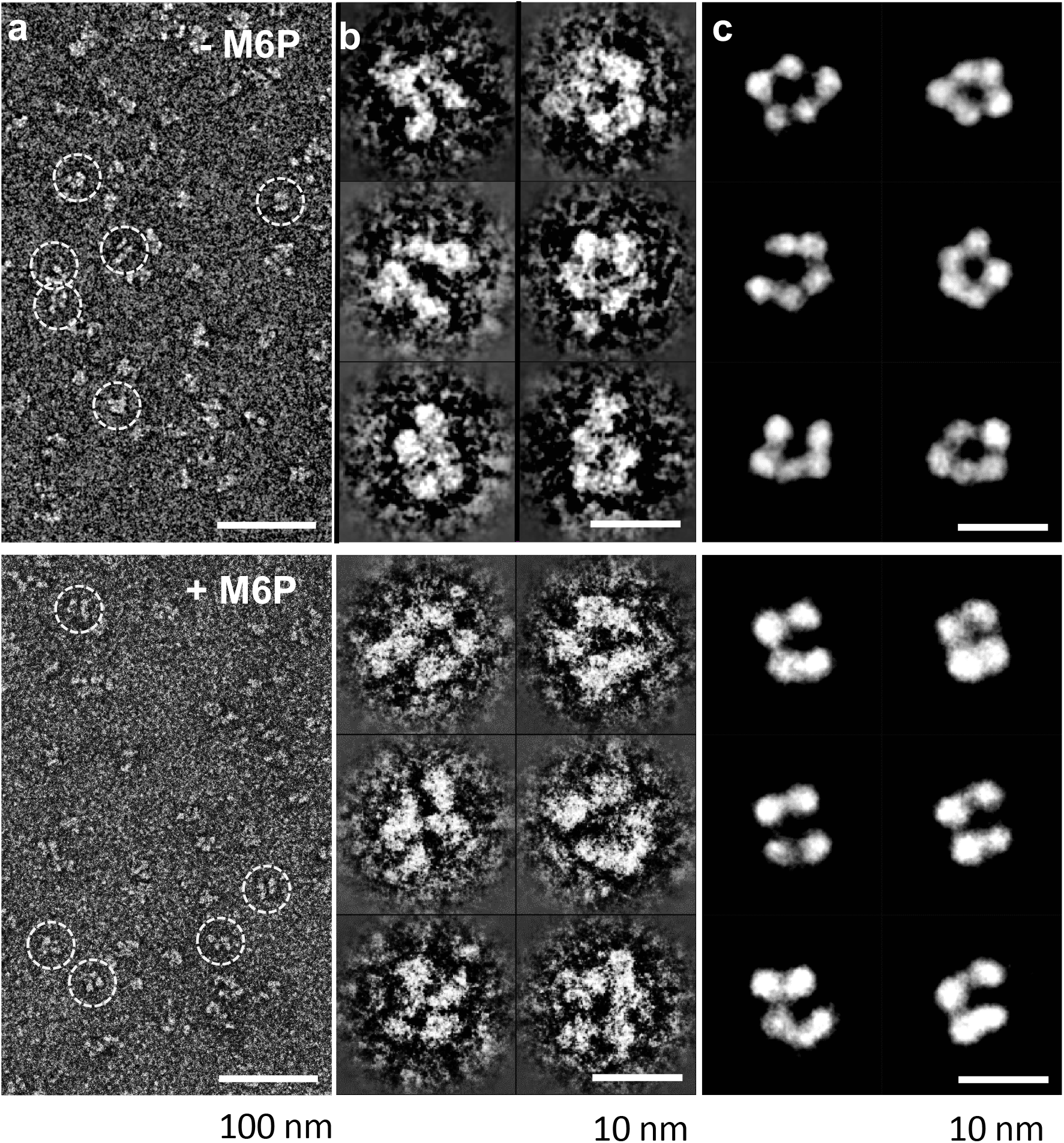
Negative stain electron microscopy of domains 1-5 of CI-MPR in the absence (upper panels) and presence (lower panels) of M6P. **a,** A survey of a negative-stain TEM image of the sample of CI-MPR domains 1-5. **b,** Six representative images of reference-free class averages of the particles of CI-MPR domains 1-5. **c,** Six representative class average images of the particles of CI-MPR domain 1-5 selected from a pool of 35 representative reference-free class averages.

### Adjacent domains are required to achieve high affinity binding

Our previous studies to identify and map the location of M6P binding sites in CI-MPR show that individual CRDs, while retaining their glycan specificity (phosphomonester or phosphodiester), bind ligand with a lower affinity than in the context of a construct containing multiple domains. For example, domain 3 bound the lysosomal enzyme β-glucuronidase with ~1,000-fold lower affinity than domains 1-3 (K_*D*_ = 500 nM versus 0.5 nM) ^42,43^, and our crystal structure of domains 1-3 (PDB 1SYO) indicate interdomain interactions, particularly with domain 1, stabilize the binding pocket of domain 3^26,31^. In contrast, the two domains immediately C-terminal to domain 3 have a minimal effect on the phosphomonoester-specific binding activity of domain 3: surface plasmon resonance (SPR) analyses show domains 1-3 and domains 1-5 exhibit similar affinity to the lysosomal enzyme PPT1 that contains predominantly phosphomonoesters (Extended Data Fig. 4 a-d). Thus, the domains 1-3 construct is sufficient to convey high affinity binding and the presence of domains 4 and 5 are not required for proper carbohydrate binding function.

To evaluate the phosphodiester-specific binding site in domain 5, we used SPR and the lysosomal enzyme acid α-glucosidase (GAA) that has been modified to contain only phosphodiesters on its *N*-glycans^38^. The presence of the additional four N-terminal domains significantly increases the affinity of domain 5 for GAA phosphodiester (K_*D*_ = ~60 nM) (Fig. 4), which is an ~150-fold higher affinity than we previously showed for a construct encoding domain 5 alone^24^. We reported a similar finding of increased binding affinity (~60-fold) comparing a construct encoding domains 5-9 with that of domain 5 alone^24^. This change in affinity cannot be attributed to protein misfolding because our NMR solution structure^27^ demonstrates that domain 5 alone is stable and exhibits the MRH β-barrel fold found in CD-MPR and all domains of CI-MPR solved to date. As expected, domains 1-5 bound with high affinity to the lysosomal enzyme GAA, modified to contain phosphodiesters or phosphomonoester, and PPT1 (Fig. 4). Together, these data are consistent with the hypothesis that additional domain(s) contribute to the high affinity binding of domain 5 through protein-protein interactions that serve to stabilize the binding site and/or through the presence of a secondary binding site(s).

**Fig. 4.**
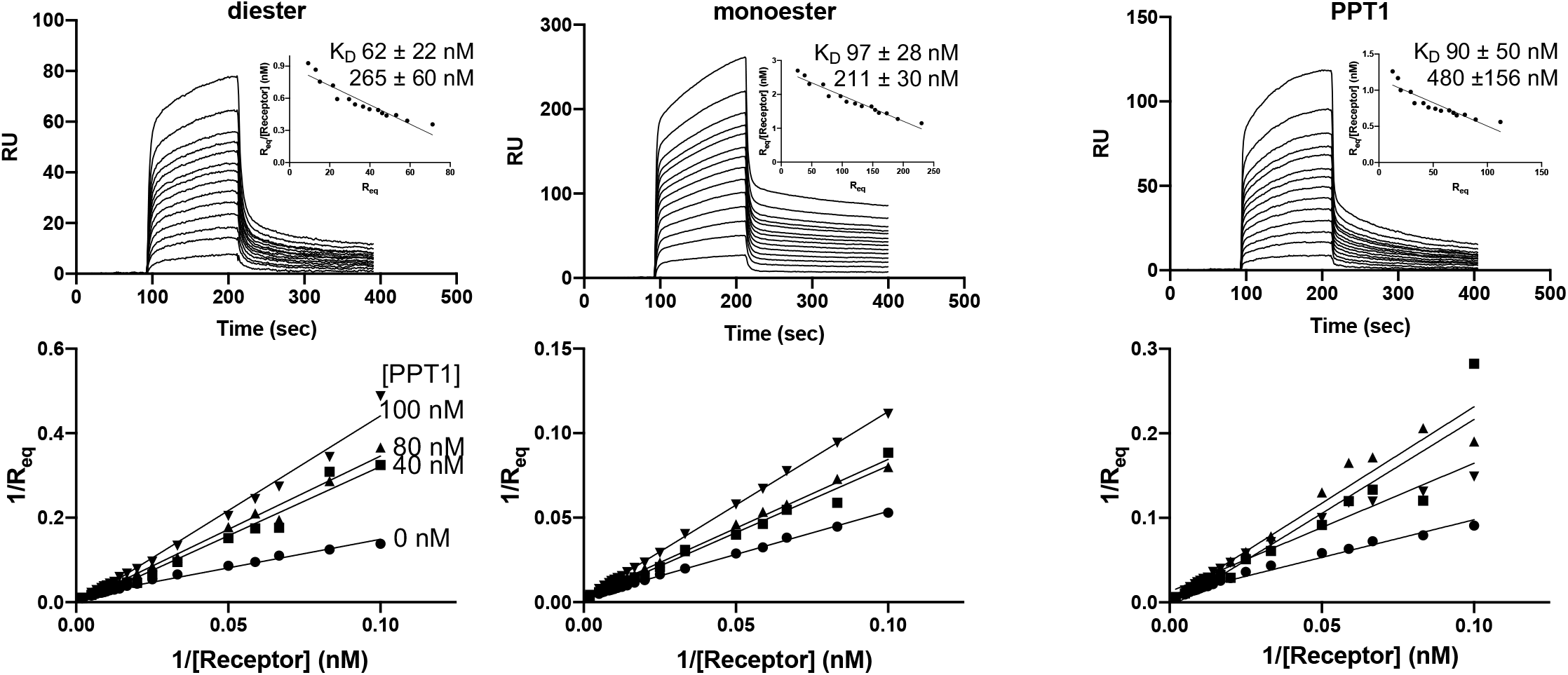
Ligand binding properties of domains 1-5 as assessed by SPR. Sensorgrams for domains 1-5 truncated protein (10 to 120 nM) flowing over GAA-phosphodiester **(upper left)** or GAA-monoester **(upper center)** and PPT1 **(upper right)** surfaces. Accompanying competitive inhibition plots (**lower panels**) for 10 nM to 120 nM domains 1-5 with 0-100 nM PPT1 as indicated in the lower left panel. Upper panel inset graphs show Scatchard plots of data.

We next evaluated if carbohydrate binding to one CRD (domain 3) affects ligand binding to the second CRD (domain 5). For these SPR studies, increasing concentrations of domains 1-5 was pre-incubated with either buffer or a fixed amount of PPT1 (has phosphomonoester *N*-glycans, see Extended Data Fig.4b) before flowing over a sensor chip immobilized with GAA phosphodiester, GAA phosphomonoester, or PPT1. The resulting sensorgrams were analyzed and displayed by double reciprocal plots (Fig. 4, lower panel). Non-parallel concentration curves intersecting near the origin were obtained for the phosphomonoester surfaces (GAA phosphomonoester and PPT1), showing that as expected PPT1 competitively competes against phosphomonoester ligand binding by domains 1-5. When the receptor-PPT1 complex (PPT1 pre-bound to domain 3 leaving domain 5 unbound) was flowed over a GAA phosphodiester surface, similar results to those obtained for the phosphomonoester surfaces were observed (Fig. 4): PPT1 is able to inhibit phosphodiester binding. One explanation is that PPT1 binds to domain 3 and sterically blocks domain 5 from binding GAA phosphodiester. However, this possibility seems unlikely based on SAXS data: we observed the envelope for PPT1 (bound to domain 3) is elongated and points away from the N-terminus and the rest of the receptor (Fig. 2c), showing that PPT1 is not in contact with domain 5. Another possibility is that PPT1 binding to domain 3 causes a rearrangement of domains such that the binding site of domain 5 is no longer accessible by ligand. This latter possibility is consistent with our crystal structures that illustrate domain interactions can be dramatically altered, such as between domain 3 and domains 1 and 2 (Fig. 1h). This later hypothesis is further evaluated by protein footprinting studies (see below).

### Mapping PPT1 interactions

To further interrogate receptor-PPT1 interactions, we turned to hydroxyl radical protein footprinting (HRPF) by means of fast photochemical oxidation of proteins (FPOP)^44^ as a method to compare protein topography between two structural states (*e.g*., ligand-bound versus ligand-free). Briefly, proteins are allowed to react with a high concentration of very short-lived hydroxyl radicals generated *in situ*. These hydroxyl radicals diffuse to the surface of the protein, where they oxidize amino acid side chains forming stable protein oxidation products at the site of oxidation. The rate of this oxidation reaction is directly proportional to the solvent accessible surface area of the amino acid. Changes in amino acid accessibility at the protein surface can be localized and measured by monitoring the rate of reaction of these surface amino acids: occlusion of that portion of the protein surface results in a decrease in the apparent rate of oxidation, while exposure of that portion of the protein surface results in an increase in the rate of oxidation^45^. These stable oxidation products are then measured through liquid chromatography-tandem mass spectrometry (LC-MS/MS)^46^. FPOP analysis of the changes in topography of CI-MPR domains 1-5 upon addition of PPT1 reveals three regions of change distributed over 4 of the 5 domains (Fig. 5a and Extended Data Fig. 5a). PPT1 binding: 1) occludes highly species conserved (Extended Data Fig. 2) regions of the interface between domains 1 and 3, including loop C (previously identified as being part of the high-affinity carbohydrate binding site) (Fig. 5b)^31^; 2) occludes C-terminal region of front β-sheet (strands 2-4) of domain 5 (Fig. 5c); 3) causes domain 4 to exhibit a topographical rearrangement resulting in occlusion of some surfaces and exposure of others (Fig. 5d, Extended Data Fig. b-d). Together, these findings demonstrate a binding-induced conformational change of CI-MPR. Because the interaction of PPT1 with domains 1-5 occurs with a 1:1 stoichiometry (Extended Data Fig. 3c) that is supported by SPR analyses showing only one CRD can be engaged in ligand binding at a time (Fig. 4) these data are consistent with PPT1’s phosphomonoester-containing *N*-glycans engaging domain 3 to cause a reorientation of domains such that domain 5 is no longer able to bind ligand (Extended Data Fig. 5d).

**Fig. 5.**
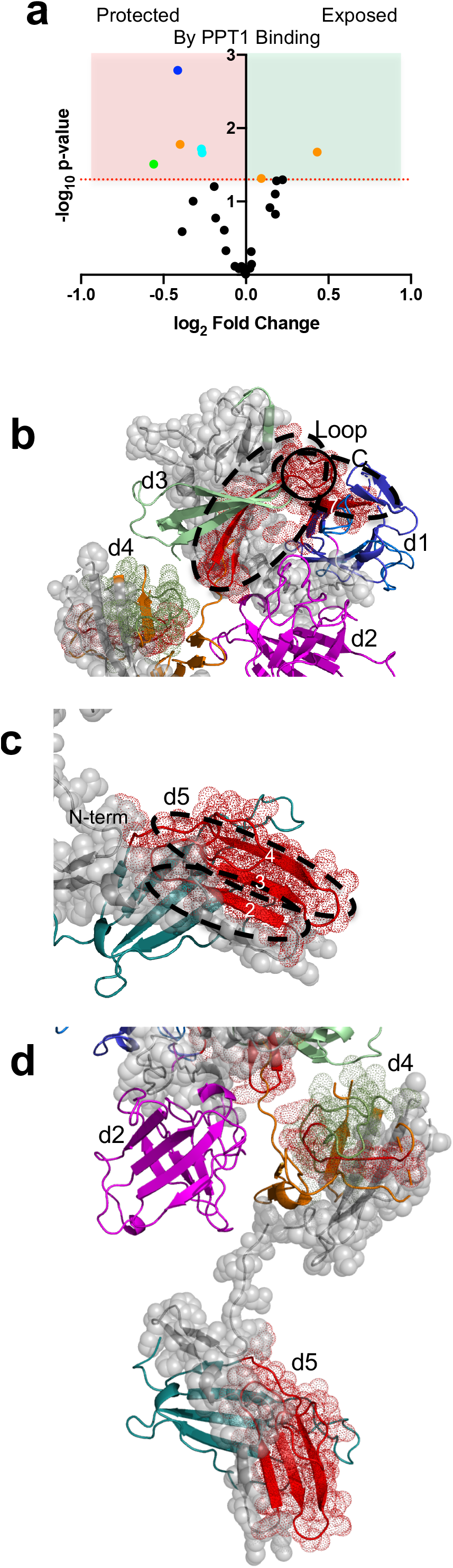
FPOP analysis of domains 1-5 in the absence and presence of PPT1 at pH 6.5. **a,** FPOP comparison of domains 1-5 alone and in the presence of PPT1 reveals peptides protected (red shaded region and colored by domain: d1, blue; d2, magenta; d3, green; d4, orange; or d5, cyan) or exposed (green shaded region and colored by domain) upon binding, while most peptides (black circles) show no statistically significant (p≤0.05) changes (unshaded region). **b,** Comparison of peptides (mapped onto SAXS generated model of bound domain 3 and unbound domain 5) with regard to changes in oxidation in the absence and presence of PPT1 ligand (red mesh and ribbon, peptides more protected, green mesh and ribbon, peptides less protected in presence of PPT1, grey spheres, no data available). **c,** Domain 5 peptides showing protection from oxidation in the presence of PPT1 mapped onto model in **(b)**. **d,** Peptides in domains 4 and 5 showing changes in exposure and protection in the presence of PPT1 mapped onto current model of domain 5 being ligand free and domain 3 ligand bound.

### Evidence for a secondary carbohydrate binding site

Two peptides in domain 3 experience decreased rates of oxidation upon the addition of PPT1, indicating a decrease in their solvent accessibility (Extended Data Fig. 5e-f). Peptide 370-391 is located in the M6P binding site of domain 3 and contains the previously identified residue R391 that is highly conserved and is essential for high affinity M6P binding^43^. The observed decreased oxidation rate of peptide 370-391 is consistent with PPT1 binding to this region of the receptor and altering solvent accessibility to hydroxyl radicals. The second peptide, 91-101, is located on β-strand 7 on the C-terminal β sheet of domain 1. As shown in our crystal structures of domains 1-3 bound to carbohydrate, domain 1’s β-strand 7 is positioned across from domain 3’s M6P binding site (Extended Data Fig. 5e-f). Although this peptide in domain 1 is outside the known M6P binding region in domain 3, its close proximity coupled with its altered oxidation rate upon PPT1 binding raises the possibility it functions as part of a secondary site of carbohydrate interaction. The existence of a secondary, lower affinity binding site is consistent with SPR analyses that showed high and low binding affinities for CI-MPR domains 1-3 upon PPT1 binding (Fig. 4, upper insets).

*In silico* approaches were used to further explore this possibility of a secondary binding site independent of a known CRD. Initial docking experiments were conducted on the modified (bound domain 3, unbound domain 5) crystal structure of domains 1-5 bound using the FTMap server to identify possible small molecule interaction sites (Fig. 6a)^47^. One of the identified ‘hotspots’ overlaps with the species conserved region of peptide 91-101 of domain 1’s β-strand 7 identified by HRPF (Extended Data Fig. 5e-f). Bidentate binding to two arms of a single oligosaccharide was ruled out due to previously reported findings by Yamaguchi *et al.* who demonstrated that a cyclic glycopeptide had a more than 10-fold higher affinity for CI-MPR than a phosphoglycoprotein ligand harboring a single M6P-phosphoglycan and that linearization of the cyclic glycopeptide through reduction of the disulfide resulted in an ~20-fold loss in affinity^48^. However, inspection of the X-ray structure (PDB 1EI9) revealed that PPT1’s three *N*-glycans were grouped in close proximity to each other and molecular dynamics simulations demonstrate the inherent flexibility of glycans (Fig 6b). Lyly *et al*. had previously shown the glycan on N232 is critical for proper trafficking of PPT1 to the lysosome^49^. Docking of the glycan on N232 into the binding site of domain 3 allows the glycan of N212 to be in close proximity to the ‘hot spot’ described above, near strand 7 of domain 1, introducing a low affinity interaction site with domain 1 in a region that includes its β-strand 7 (Fig. 6c). Lysosomal enzymes typically possess multiple glycosylation sites, like PPT1. Some via oligomerization, like the tetrameric β-glucuronidase with four *N*-glycans per monomer^50^, present a more expansive spatial array of phosphorylated glycans that together can enhance the engagement with CI-MPR’s CRDs to obtain high affinity binding via multivalent interactions. The existence of low affinity binding sites, such as that proposed above in domain 1, would enhance the avidity of these interactions.

**Fig. 6.**
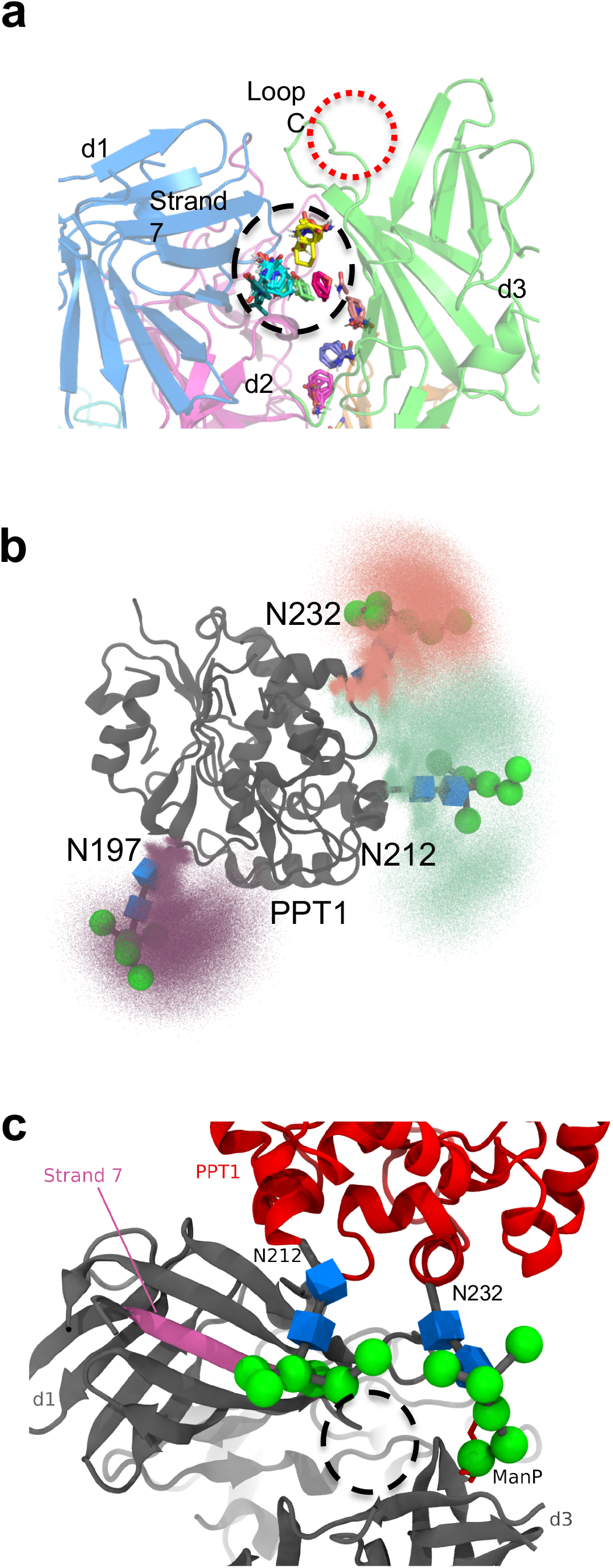
Possible secondary site of oligosaccharide interaction. **a**, Small molecule ‘hot spots’ identified through the use of the FTMap server (http://ftmap.bu.edu/login.php) are shown on the model of domains 1-5 bound to ligand in domain 3. Region near secondary site proposed from analysis of FPOP data is circled in black. Domains are labeled and colored as previously described. Loop C of the M6P binding site (circled in red) is labeled. **b**, Molecular dynamics simulations were used to map the extent of movement the glycans of PPT1 are able to undergo. **c,** PPT1 (PDB 1EI9) (red) structure is overlaid on the model of domains 1-5 (dark grey) with domain 3 bound and domain 5 in the unbound position with the oligosaccharide on N232 resting near Loop C (binding site) of domain 3 and the oligosaccharide on N212 located near strand 7 (magenta) of domain 1. Small molecule ‘hot spots’ are circle in black line.

### CI-MPR undergoes domain rearrangement at pH of late endosome

Recognition and binding of lysosomal enzymes by CI-MPR represents only part of this receptor’s function. The role of this receptor in populating the lysosome with acid hydrolases is not complete until the receptor releases its cargo in the acidic pH environment of the endosome (~pH 5). This release process is critical since neutralization of intracellular compartments results in excessive secretion of lysosomal enzymes, with MPRs being ‘trapped’ to their cargo^51^. Size exclusion chromatography (SEC) of three constructs (domains 1-15, domains 1-5 and domains 7-15) shows that as the pH becomes more acidic, CI-MPR exhibits a more compact Stokes radius (Fig. 7a), and these changes are not confined to a particular region of the receptor. However, when the change in the calculated Stokes radius for each construct is normalized per number of domains in the construct, the N-terminal 5 domains undergo the largest change in radius: they compact on each other the most (Fig. 7b). We again turned to HRPF to evaluate conformational changes of CI-MPR as a consequence of pH. These analyses show widespread changes in conformation of domains 1-5 upon a change in pH from 6.5 to 4.5, with the changes more extensive than that observed upon the addition of PPT1 (Fig. 7c, Extended Data Fig. 6 and Fig. 5a). For example, domain 2, which had no residues with significant changes in susceptibility to hydroxy radical treatment upon PPT1 addition, showed large regions of protection from oxidation when altering the pH from 6.5 to 4.5. The increase in the regions of protection from oxidation upon acidification is consistent with SEC data and shows that CI-MPR undergoes significant domain rearrangement and a compaction of its overall conformation. Additionally, the two peptides (91-101 and 370-391) previously observed gaining protection from oxidation in the presence of PPT1 report higher degrees of modification in transitioning to acidic pH, a finding consistent with ligand release in the acidic environment of endosomes.

**Fig. 7.**
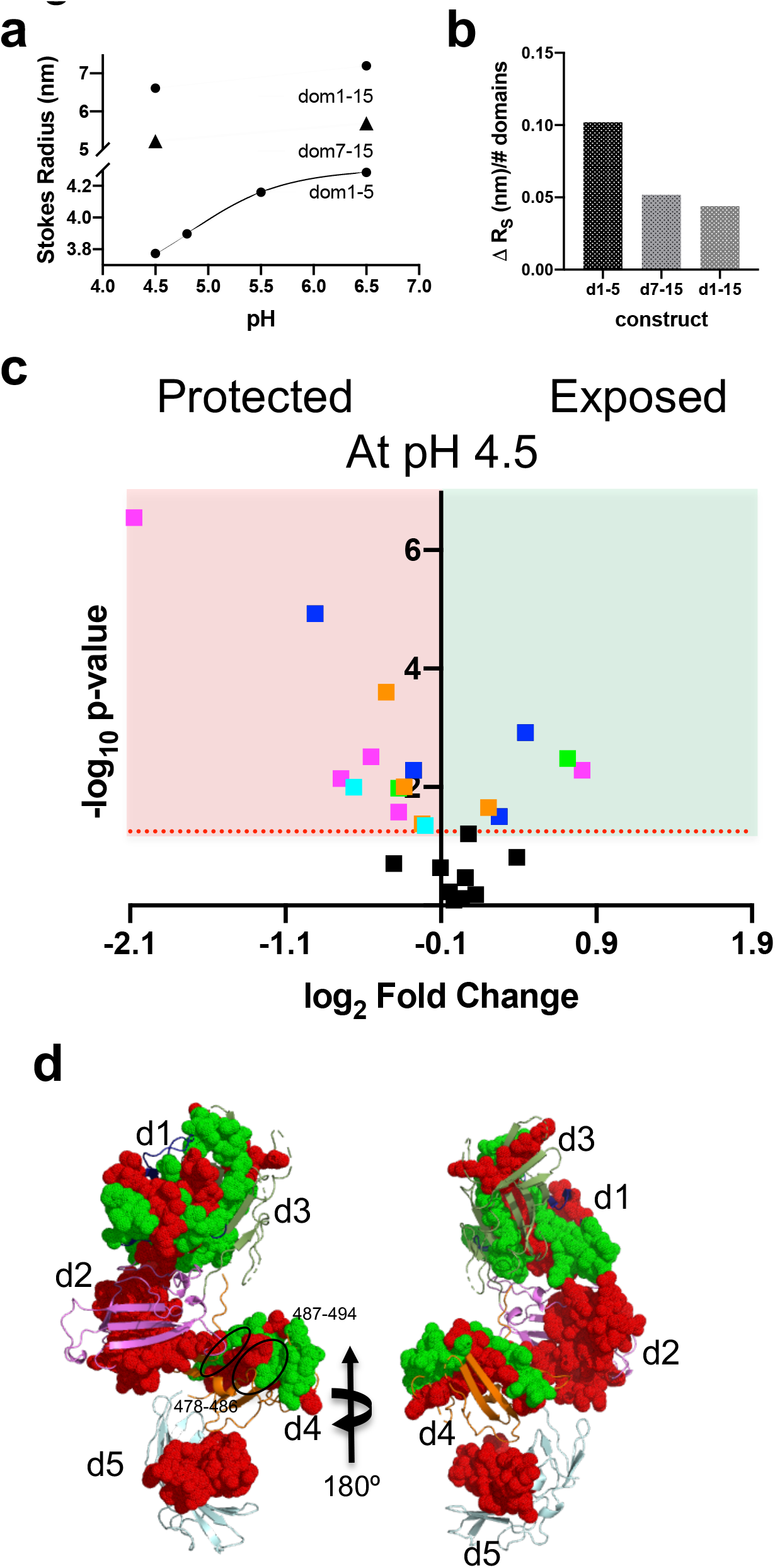
Domain 1-5 adopts a more compact conformation at pH 4.5 compared to pH 6.5. **a,** Plot of changes in calculated Stokes radius of domains 1-5, domains 7-15 and domains 1-15 with changes of pH. **b,** Changes in calculated Stokes radius for domains 1-5, 7-15 and 1-15 normalized per number of domains in each construct. **c**, FPOP analysis of domains 1-5 at pH 6.5 versus 4.5 reveals peptides protected (red shaded region and colored by domain: d1, blue; d2, magenta; d3, green; d4, orange; or d5, cyan) or exposed (green shaded region and colored by domain) upon lowering of pH to that of endosome. Overall more peptides show statistically significant (p≤0.05) changes compared to PPT1 binding (Fig. 5a), while fewer peptides (black boxes) show no statistically significant changes (unshaded region). **d**, Peptides showing a greater degree of protection at pH 4.5 versus 6.5 are mapped as red spheres onto the SAXS based model of domains 1-5 in the absence of ligands, while those showing less protection are mapped as green spheres onto the model. Model regions undergoing no statistically significant changes or lacking data are represented as ribbons.

### Conclusions and Future Directions

Our current and past X-ray crystal structures, coupled with SAXS, FPOP and SPR analyses, allows us for the first time to propose an allosteric mechanism for the functioning of CI-MPR: ligand binding to one site influences binding to a second. Carbohydrate binding to either domain 3 or 5 facilitates a change in domain orientation, thereby blocking ligand accessibility and/or stability of the binding pocket. Despite crystallization in the presence of M6P, no ligand was detected in domain 3. This finding can be explained by a *N*-glycan of a crystallographic neighbor occupying the binding pocket of domain 5, causing in a dramatic change in domain 3’s orientation. This change results in the loss of key, stabilizing contacts between the M6P binding pocket of domain 3 and residues in domains 1 and 2. Although carbohydrate specificity is contained within the individual CRD, high affinity binding requires the presence of additional domains, as we reported for domain 3^31^ and domain 5 (in this report and see^24^). Additionally, domain 15 exhibits enhanced binding affinity (~85-fold) to a lysosomal enzyme in the presence of domain 14^25^. This common theme of allosteric regulation of ligand binding extends to the non-M6P-containing peptide, IGF2, in which contacts with domain 13 enhances the binding affinity of domain 11 for IGF2 by ~10-fold^52^. Taken together, these data lead us to propose a new model of carbohydrate binding by domains 1-5 of CI-MPR whereby a second, lower affinity site on neighboring domain is occupied by an adjacent oligosaccharide of the lysosomal enzyme. We report the first structural view of a complex between CI-MPR and a lysosomal enzyme, PPT1, providing insight for future studies into developing therapy for newborns with infantile neuronal ceroid lipofuscinosis who are deficient in this enzyme. Recent studies showing PPT1-dependent depalmitoylation stabilizes the lysosomal localization of v-ATPase subunits, which directly impacts lysosomal acidification required for autophagy-lysosomal function and mTOR signaling, further emphasizes the need for proper MPR-mediated delivery of PPT1 to the lysosome^53^.

Several questions remain to be answered in order to fully understand how ligand binding regulates CI-MPR structure and function. For example, does lysosomal enzyme binding elicit allosteric effects on CI-MPR’s non-M6P-containing ligands, IGF2, plasminogen, and/or uPAR? Conversely, do these non-M6P-containing ligands modify carbohydrate binding activity of one, several, or all four of CI-MPR’s CRDs? Are the conformational dynamics of the N-terminal five domains impacted by the receptor’s C-terminal 10 domains? With respect to pH dependent release of ligand, we made an interesting observation while purifying our constructs over a PPT1 affinity column: unlike domains 7-15, constructs containing the N-terminal domains 1-3 or 1-5, along with domains 1-15, eluted efficiently from the column upon reducing the buffer pH to 4.5 (Extended Data Fig. 7). Similarly, we found that domains 1-3, but not domains 7-9 or 7-11, eluted from a pentamannosyl phosphate-agarose column at pH 4.6^54^. Consistent with our SEC data showing a compact conformation upon acidification (Fig. 7), we propose that CI-MPR adopts a bent conformation as it traffics to acidified endosomal compartments such that its N-terminal region interacts with C-terminal domain(s) to trigger ligand release by the CRDs in domains 9 and/or 15. Taken together, it is intriguing to speculate that the N-terminal 5 domains play a predominant role in regulating CI-MPR’s ability to bind and release its diverse ligands. In conclusion, CI-MPR function appears to be tightly regulated by allosterism and future studies are needed to further understand domain interplay of this conformationally dynamic receptor.

## METHODS

### Generation and expression of human CI-MPR constructs

DNA sequences corresponding to domains 1-3 (residues 1-433), 1-5 (residues 1-726), domains 7-15 (residues 888-2296) and domains 1-15 (residues 1-2296) (numbering does not include the N-terminal signal sequence (residues 1-35) of the human CI-MPR were amplified directly from the human clone (GeneBank Accession No. J03528) obtained from the American Type Culture Collection (pGEM-8 ATCC 95661) using standard polymerase chain reaction (PCR) methods. Mutant cDNAs were generated using DpnI mediated site-directed mutagenesis and confirmed by DNA sequencing. All constructs were cloned into pVL1392 modified to contain the native bovine CI-MPR N-terminal signal sequence followed by a *NotI* sequence and a C-terminal thrombin, hexahistidine, an Avi tag and *XbaI* and used to generate Baculovirus using the BestBac method (Expression Systems). *Spodoptera frugiperda* (Sf9) cells were infected with baculovirus at a density of 3.0 × 10^6^ cells per ml for 96 hours at 27° C in ESF 991 with 1% Production Boost Additive added after 24 hours. Cells were removed from the medium by centrifugation at 1,000*g.*

### Purification, crystallization and data collection of human CI-MPR domains1-5 protein at pH 5.5

Medium was concentrated to ~40ml using Amicon stir cells prior to dialysis against 3 times 4 liters of 20 mM Tris, pH 7.5 at 22° C, 150 mM NaCl. Protein solutions were centrifuged at 20,000g to remove particulate matter prior to loading on 5 ml Ni-NTA columns. Resin was washed with 20 mM Tris pH 7.6, 300 mM NaCl, and 20 mM imidazole before elution with 20 mM Tris pH 7.6, 300 mM NaCl, and 100 mM imidazole. Fractions were analyzed by SDS-Page, pooled and concentrated to 1 mg/ml before overnight dialysis at 4° C into 20 mM Tris, pH 7.6 at 22° C, 150 mM NaCl. C-terminal tags were removed by incubation overnight at 4° C with Thrombin (Sigma). Thrombin was removed by passage over benzamidine agarose beads. Protein was incubated overnight with pNGasF to remove N-linked glyans followed by passage over Ni-NTA agarose to remove His-tagged PNGaseF. Domains 1-5 protein was then passed over a 10/300 Superdex G200 column equilibrated in 20 mM Tris pH 7.4 at 22° C, 150 mM NaCl to remove any remaining aggregates or contaminants. Protein was concentrated to 7 mg/ml and incubated with 10 mM M6P and 10 mM MnCl_2_. Initial crystallization hits were found with Molecular dimensions JCSG-plus and optimized to 100 mM ammonium citrate dibasic, pH 5.5, 20% PEG 3350. Crystals were cryoprotected in reservoir solution + 25% glycerol and frozen in liquid nitrogen. Data was collected on using a Rigaku M007 detector equipped with Osmic mirrors and an R-AXIS IV++ detector. Data was processed and scaled with HKL2000 software^55^.

### Purification, crystallization, data collection and processing of human CI-MPR domains 1-5 at pH 7.0

Protein was prepared as described above except without PNGaseF digestion and additional purification over a 5 ml pentamannosyl phosphate agarose affinity column^56^ equilibrated in 50 mM imidazole, pH 6.5, 150 mM NaCl, 5 m β-glycerol phosphate. Protein was eluted by the addition of 10 mM M6P to column buffer. Protein was concentrated to 7 mg/ml and incubated with 10 mM MnCl_2_. A crystallization hit was identified using Molecular dimensions JCSG-plus (100 mM HEPES, 30% Jeffamine ED-2003. Crystals were cryoprotected in reservoir solution + 20% glycerol and frozen in liquid nitrogen. Data was collected at APS beamline 17-ID (IMCA-CAT) at 100° K, processed using autoPROC^57^.

### Structure Determination of human CI-MPR domains 1-5

Phases for the pH 5.5 conditions structure were determined by Phaser in CCP4i^58^ using homology models generated using the Swiss-Model server. The model was iteratively refined using PHENIX^59^ and manually rebuild in COOT^60,61^ and showed reasonable stereochemistry, with 86.7% in the Ramachandran favored zones, 10.7% in the allowed and 2.5% outliers. The pH 7.0 structure was also solved by MR using PHASER and the domain 1-5 structure previously solved. The model was refined using PHENIX and iteratively rebuild in COOT and showed reasonable stereochemistry, with 88.4% in the Ramachandran favored zones, 10.0% in the allowed and 1.6 % outliers.

### Expression and purification of PPT1

Recombinant human PPT1 protein was grown and purified following the protocol outlined in Lu *et al.*, 2010^62^.

### Purification of human CI-MPR domain1-5 protein for SAXS, SPR, negative stain, EM and SEC

Cells were removed from the medium by centrifugation at 1,000*g* and the pH of the medium was adjusted to pH 8.0 with 10M NaOH. Precipitates were removed from the medium by centrifugation at 4,000*g* before loading over 5 ml of PROTEINDEX Ni-Penta agarose 6 fast flow resin (Marvelgent Biosciences). Resin was washed with 20 mM Tris pH 7.6, 300 mM NaCl, and 20 mM imidazole before elution with 20 mM Tris pH 7.6, 300 mM NaCl, and 100 mM imidazole. Fractions were analyzed by SDS-Page, pooled and concentrated to 1 mg/ml before overnight dialysis at 4° C into 20 mM Tris, pH 7.6 at 22° C, 150 mM NaCl. C-terminal tags were removed by incubation overnight at 4° C with Thrombin (Sigma). Thrombin was removed by passage over benzamidine agarose beads. Domain 1-5 protein was also incubated with PNGaseF overnight at 22° C to remove N-linked glycans. Proteins were passed over Ni-NTA agarose to remove cleaved His-tags as well as PNGaseF (His-tagged). Flow through and proteins eluted with 20 mM Tris pH 7.5 at 22° C, 150 mM NaCl were combined and dialyzed overnight at 22° C in column buffer (50 mM imidazole, pH 6.5, 150 mM NaCl, 5 m β-glycerol phosphate, 10 mM MnCl_2_). Protein was further purified by passing dialyzed proteins over a 1 ml PPT1 amine coupled agarose resin column (10 mg/ml) (Pierce NHS-activated agarose slurry). Loaded protein was washed with column buffer prior to elution with 20 mM MES, pH 4.5, 150 mM NaCl, 5 m β-glycerol phosphate, 10 mM MnCl_2_ and neutralization with Tris buffer to pH 7.4. Any aggregates or contaminants were removed by passaged over a Superdex G200 10/300 column equilibrated in column buffer without MnCl_2_. Monomeric fractions were pooled and stored at 4° C. Protein concentration was determined using the Bradford assay (Bio-Rad) with bovine serum albumin as the standard.

### Purification of CI-MPR domains 1-3, 7-15 and domains 1-15 for SPR and/or SEC

Proteins were expressed in Sf9 cells grown, harvested and purified as described in the previous section without treatment with thrombin and PNGaseF.

### LC-MS/MS analysis of PPT1 glycosylation

Aliqouts of PPT1 (20 μg) were reduced, carboxyamidomethylated, dialyzed against nanopure water at 4°C overnight, and then dried in a Speed Vac. The dried, desalted sample was resuspended and digested with trypsin (Promega, sequence grade) at 37 °C overnight. Following digestion, the sample was again dried and subsequently resuspended in solvent A (0.1% formic acid in water) and passed through a 0.2 μm filter (Nanosep, PALL) prior to analysis by liquid chromatography-tandem mass spectrometric analysis (LC-MS/MS).

LC-MS/MS analysis was performed on an Orbitrap-Fusion equipped with an EASY nanospray source and Ultimate3000 autosampler LC system (Thermo Fisher). Resuspended tryptic peptides were chromatographed on a nano-C18 column (Acclaim pepMap RSLC, 75 μm × 150 mm, C18, 2 μm) with an 80-min gradient of increasing mobile phase B (80% acetonitrile, 0.1% formic acid in distilled H2O) at a flow rate of 300 nl/min routed directly into the mass spectrometer. Full MS spectra were collected at 60,000 resolution in FT mode and MS/MS spectra were obtained for each precursor ion by data-dependent scans (top-speed scan, 3 sec) utilizing CID, HCD, or ETD activation and subsequent detection in FT mode.

Phosphorylated glycopeptides were annotated by manual data interpretation of the LC-MS/MS data following initial processing by Byonic software (Protein Metrics). Byonic parameters were set to allow 20 ppm of precursor ion monoisotopic mass tolerance and 20 ppm of fragment ion tolerance. Byonic searches were performed against the human palmitoyl-protein thioesterase 1 (PPT1) sequence allowing modification with phosphorylated and non-phosphorylated human/mammalian N-glycans.

### Surface Plasmon Resonance studies of CI-MPR domains 1-5

All SPR measurements were performed at 25° C using a Biacore 3000 instrument (BIAcore, GE Healthcare, Piscataway, NJ) as described previous^24,25^. Purified lysosomal enzymes (GAA mono- and diester, PPT1) were immobilized at a density of ~1000 RU on a CM5 sensor chip by primary amine coupling following manufacturer’s procedure. The reference surface was treated the same way except for no protein addition. Purified domains 1-5 and domains 1-5 with PPT1 were prepared in 50 mM imidazole, 150 mM NaCl, 5 mM MgCl_2_, 5 mM MnCl_2_, 5 mM CaCl_2_, pH 6.5 supplemented with 0.005% (v/v) P20. All sample were incubated for 2 hours prior to loading on instrument. Samples were injected in a volume of 80 μl over the reference and coupled flow cells at a flow rate of 40 μl/min for 2 minutes prior to dissociation with buffer alone for 2 min. The sensor chip surfaces were regenerated with a 20 μl injection of 10 mM HCl at a flow rate of 10 μl/min and allowed to re-equilibrate with running buffer for 1 min. prior to the next injection. The response at equilibrium (R_eq_) was determined for each concentration of protein/complex by averaging the response over a 10 s time span within the steady state region of the sensogram (BIAevaluation software package, 4.0.1). Scatchard analysis was performed to determine linear regions (10-40 nM) and (50-200 nM). The R_eq_ was plotted for these two regions versus the concentration of protein and fit to a 1:1 binding isotherm. All responses were double-referenced by subtracting the change in refractive index for the flow cell derivatized in the absence of protein from the binding sensorgrams^63^.

### SAXS data collection on CI-MPR domains 1-5 in presence and absence of ligands

SAXS were performed at BioCAT (Sector 18) Advanced Photon Source utilizing a Pilatus 1M detector. Data was collected at ~20° C with a wavelength of 1.033 Å and ~3.5m sample-to-detector distance (*q* range = 0.00535-0.387 Å^−1^). Prior to introduction into the stationary SAXS quartz capillary (1.5mm ID, 1.52 mm OD), 0.5 mg of domains 1-5 protein was incubated with10 mM M6P in 50 mM imidazole, pH 6.5, 150 mM NaCl, 5 mM β-glycerolphosphate for 1 hour at 22° C. Batch mode SAXS data were collected on hd1-5 alone, and in complex with M6P. SEC-SAXS was performed on domains 1-5 in complex with PPT1, PPT1 along and domains 1-5 protein alone. For the complex, 1.25mg of domains 1-5 with 5 mg of PPT1 in above imidazole for for 1 hr at 22° C prior to data collection. SEC/SAXS data was collected simultaneously (0.5s exposures collected every 3 s) upon elution from a 10/300 Superdex G200 Increase column equilibrated in matched buffer and at a flow rate of 0.75 ml/min. Free domains 1-5 protein could not be completely separated from that in complex with PPT1 as seen in chromatogram. Exposures flanking the elution peaks were averaged to generate the I(q) vs. q curve for the buffer and then subtracted from the elution peak curves to obtain the sample SAXS curves. Data were processed with Primus^64^ and *Ab initio* dummy atom modeling was done with DAMMIF^65^. The merged SAXS curves were used to generate pair distribution functions, P(r), and Kratky plots (PRIMUS). The flexibility analysis curves were generated using SCATTER 3.0 software. The FoXS server was used to compute the SAXS profile using the coordinates from the pH 5.5 structure (PDB 6P8I). The MultiFoXS server was used to calculate the population-weighted ensemble fitting to the unbound protein scattering curves.

### Electron Microscopy (EM) on CI-MPR domains 1-5 in the presence and absence of M6P

The negative stained (NS) EM specimens of domains 1-5 and the domains 1-5 bound to M6P were prepared as described above. In brief, the samples were diluted to ~0.001 μg mL^−1^ with sample buffer. An aliquot (approximately 4 μL) of diluted sample was placed on an ultra-thin carbon-coated 200-mesh copper grid (CF200-Cu-UL, Electron Microscopy Sciences, Hatfield, PA, USA, and Cu-200CN, Pacific Grid-Tech, San Francisco, CA, USA) that had been glow-discharged for 15 s. After 1-min incubation, the excess solution on the grid was blotted with filter paper. The grid was then washed with water and stained with 1% (w/v) uranyl formate before air-drying with nitrogen. The EM samples were examined by using a Zeiss Libra 120 Plus TEM (Carl Zeiss NTS) operated at 120 kV high tension with a 10-20 eV energy filter. The OpNS micrographs were acquired under defocus at ~0.6 μm and a dose of ~40-90 e^−^Å^−2^ using a Gatan UltraScan 4K X 4K CCD under a magnification of 80 kx (each pixel of the micrographs corresponds to 1.48 Å in specimens). The contrast transfer function (CTF) of each micrograph was examined by using *ctffind3* software^66^ and the phase and amplitude were corrected by using the “*TF CTS*” command in SPIDER^67^ software or GCTF^68^ after the X-ray speckles were removed. Particles were then selected from the micrographs by using *boxer* (EMAN software^69^). All particles were masked by using a round mask generated from SPIDER software after a Gaussian high-pass filtering. The 50 reference-free class averages of particles were obtained by using *refine2d* (EMAN software) based on ~3,000 particles windowed from ~140 micrographs.

### Size-exclusion chromatography of truncated human CI-MPR constructs

A G200-Increase 10/300 column was run at a flow rate of 0.75 ml/min and equilibrated with either pH 6.5 buffer (20 mM imidazole, 150 mM NaCl, pH 7.5), pH 5.5 buffer (20 mM sodium citrate, 150 mM NaCl, pH 5.5) pH 4.8 buffer (20 mM sodium citrate, 150 mM NaCl, pH 4.8), or pH 4.5 buffer (20 mM sodium citrate, 150 mM NaCl, pH 4.5). Domains 1-5 protein was run at all the above listed pH values while domains 7-15 and domains 1-15 were run at pH 6.5 and pH 4.5. All proteins were injected onto the column as 50 μg in 200 μl of matched pH buffer. Stokes radius were calculated as described by La Verde *et al.*, 2017^70^ using thyroglobulin (bovine thyroid), β-amylase (wweet potato), albumin (bovine albumin), carbonic anhydrase (erythrocytes), and cytochrome c (horse heart).

### FPOP of human CI-MPR domains 1-5

A final concentration of 5 μM domains 1-5 protein was incubated in the 5 mM sodium citrate buffer in the presence or absence of 5 μM PPT1 at pH 6.5 for one hour. For FPOP at pH 4.5, 5 mM sodium citrate buffer was used to incubate CI-MPR domain 1-5 for 1 hr without PPT1. FPOP was performed as described previously^71^. Briefly, 20 μl of protein sample mixture containing 1 mM adenine, 17 mM glutamine, and 100 mM hydrogen peroxide was irradiated by flow through the path of the pulsed ultraviolet laser beam from a Compex Pro 102 KrF excimer laser (Coherent, Germany). The laser fluence was calculated to be ~10.1 mJ/mm^2^/pulse. Laser repetition rate was 15 Hz. The flow rate was adjusted to 13 μL/min to ensure a 15% exclusion volume between irradiated segments. After laser illumination, each replicate was collected in a microcentrifuge tube containing 25 μl of quench mixture that contained 0.5 μg/μl H-Met-NH_2_ and 0.5 μg/μl catalase to eliminate secondary oxidants. The adenine hydroxyl radical dosimetry readings were measured at 265 nm in nanodrop (Thermo Scientific) to ensure all the samples were exposed to equivalent amounts of hydroxyl radical^72^. All FPOP experiments were performed in triplicate for statistical analysis.

After FPOP and quenching, 50 mM Tris, pH 8.0 containing 1 mM CaCl_2,_ 5 mM DTT was added to the protein samples and incubated at 95 °C for 15 minutes to denature and reduce the protein. The sample was cooled on ice, trypsin with 1:20 ratio of trypsin:protein was added and incubated at 37 °C for 12 hours with rotation. Sample digestion was stopped by adding 0.1% formic acid and the samples were analyzed on a Dionex Ultimate 3000 nano-LC system coupled to an Orbitrap Fusion Thermo Scientific (San Jose, CA). Samples were trapped on a 300 μM id X5 mm PepMap 100, 5 μm (Thermo Scientific) C18 trapping cartridge, then back-eluted onto an Acclaim PepMap 100 C18 nanocolumn (0.75 mm × 150 mm, 2 μm, Thermo Scientific). Separation of peptides on the chromatographic system was performed using a binary gradient of solvent A (0.1% formic acid in water) and solvent B (0.1% formic acid in acetonitrile) at a flow rate of 0.30 μL/min. The peptides were eluted with a gradient consisting of 2 to 10% solvent B over 4 min, increasing to 32% B over 25 min, ramped to 95 % B over 4 min, held for 4 min, and then returned to 2% B over 2 min and held for 8 min. Peptides were eluted directly into the nanospray source of an Orbitrap Fusion instrument using a conductive nanospray emitter obtained from Thermo Scientific. All the data were collected in positive ion mode. Collision-induced dissociation CID were used to fragment peptides, with an isolation width of 3 *m/z* units. The spray voltage was set to 2400 volts, and the temperature of the heated capillary was set to 300 °C. Full MS scans were acquired from m/z 350 to 2000 followed by eight subsequent MS2 CID scans on the top eight most abundant peptide ions.

Peptides from tryptic digests of CI-MPR domain 1-5 were identified using ByOnic version v2.10.5 (Protein Metrics). The search parameters included all possible major oxidation modifications^73^ as variable modifications and the enzyme specificity was set to cleave the protein after arginine and lysine residues. The peak intensities of the unoxidized peptides and their corresponding oxidation products observed in LC-MS were used to calculate the average oxidation events per peptide in the sample as previously reported^45^. Briefly, peptide level oxidation was calculated by adding the ion intensities of all the oxidized peptides multiplied by the number of oxidation events required for the mass shift (e.g., one event for +16, two events for +32) and then divided by the sum of the ion intensities of all unoxidized and oxidized peptide masses as represented by equation 1.

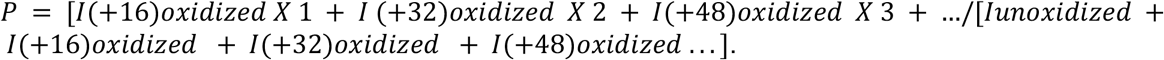

where *P* denotes the oxidation events at the peptide level and *I* values are the peak intensities of oxidized and unoxidized peptides^72^.

### The generation and energy minimization of the glycosylated structure of PPT1

The 3D structure of PPT1 (PDB code 3GRO) with M6GN2 (DManpα1-6[DManpα1-3]DManpα1-6[DManpα1-2DManpα1-3]DManpα1-4DGlcpNAcβ1-4DGlcpNAcβ1-) conjugated to N197, N212, and N232) using the glycoprotein builder available at GLYCAM-Web (www.glycam-web.org) and an in-house program that adjusts the glycosidic linkages to relieve any atomic overlaps between the conjugated glycan and the underlying protein. dsxThe glycosylated PPT1 structure was placed in a periodic box of approximately 15,000 TIP5P waters with a 10 Å buffer between the glycoprotein and the box edge. Energy minimization of all atoms was performed for 20,000 steps (10,000 steepest decent, followed by 10,000 conjugant gradient).

#### Molecular dynamics (MD) simulations

All MD simulations were performed with the CUDA implementation of the PMEMD ^74,75^ simulation code, as present in the Amber14 software suite ^76^. The GLYCAM06j force field ^77^ and Amber14SB force field ^78^were employed for the carbohydrate and protein moieties, respectively. A Berendsen barostat with a time constant of 1 ps was employed for pressure regulation, while a Langevin thermostat with a collision frequency of 2 ps^−1^ was employed for temperature regulation. A nonbonded interaction cutoff of 8 Å was employed. Long-range electrostatics were treated with the particle-mesh Ewald (PME) method ^79^. Covalent bonds involving hydrogen were constrained with the SHAKE algorithm, allowing an integration time step of 2 fs ^80^ to be employed. All simulations were performed under nPT conditions and the restraints employed were 5 kcal/mol-Å2 Cartesian. The energy minimized coordinates were equilibrated at 300K over 400 ps with restraints on the solute heavy atoms. The system was then equilibrated with restraints on the Cα atoms of the protein for 1ns, prior to performing a production MD simulation for 500 ns, but with restraints applied only to the Cα atoms of residues on either sides of gaps, namely (11, 12, 20, 21, 25, 26, 44, 45, 78, 79, 98, 99, 184, 185, 190, 191, 236 and 237. AMBER numbering).

### Modelling of multimeric interaction between PPT1 and domains 1-5

The glycosylated PPT1 structure was aligned to the Man6P in the d3 binding site of the “bound” form of the d1-5 structure. The alignment was performed by superimposing the non-reducing terminal mannose residue on the DManpα1-2DManpα1-3 arm of M6GN2 at N197 onto the complexed Man6P. This process was repeated for 100 snapshots taken at regular intervals from the MD simulation trajectory. An in-house program was employed to adjust the glycosidic linkages in N232 and N212 in order to bring N212 into contact with the region identified by the “hot-spot” analysis. The program adjusts the glycosidic linkages within known low-energy ranges^81^ while avoiding atomic overlaps, as described previously^82,83^.

### The fitting of model of domains 1-5 bound to PPT1 to SAX data

The UCSF-Chimera 1.12^84^ program was employed to fit the co-complex of d1-5 with glycosylated PPT1. Snapshots from the MD simulation of PPT1 was aligned to the “bound” form of d1-5 via N232 as described in the previous “*Modelling of multimeric interaction”* section. These structures were then fit into the SAX data using the map fitting feature of UCSF Chimera with 50 fittings per structure and a 90% cut-off.

## Data Availability

All data generated or analyzed during this study are included in the published article and its Supplemental Information. The X-ray crystal structures and structure factors of human domains 1-5 of CI-MPR at pH 5.5 and 7.0 have been deposited in Protein Data Bank under accession codes PDB 6P8I and PDB 6V02. SAXS data has been deposited at SASDBD with codes: SASDHL4 (N-terminal domains 1-5 of the cation-independent mannose-6-phosphate receptor (CI-MPR)), SASDHM4 (N-terminal domains 1-5 of the cation-independent mannose-6-phosphate receptor (CI-MPR) from SEC-SAXS), SASDHN4 (N-terminal 5 domains of the cation-independent mannose-6-phosphate receptor (CI-MPR) bound to mannose 6-phosphate (M6P)), SASDHP4 (Palmitoyl-protein thioesterase 1 (PPT1)), and SASDQ4 (N-terminal domains 1-5 of the cation-independent mannose-6-phosphate receptor (CI-MPR) in complex with palmitoyl-protein thioesterase 1 (PPT1)).

## Acknowledgements

We would like to thank Dr. Jui-Yun Lu and Dr. Sandra L. Hofmann for providing the CHO cell line expressing recombinant PPT1. We would like to thank Dr. Srinivas Chakravarthy, BioCAT, Argonne National Laboratory, sector 18 for SAXS data collection and processing during their fall 2017 SAXS data collection workshop, which was supported by grant to Thomas Irving 9 P41 GM103622 from the National Institute of General Medical Sciences of the National Institutes of Health (NIH).”. We would like to thank Richard Bohnsack for generating the human CI-MPR domains 1-5 clone and Dr. Lei Zhang’s early screening of the sample by negative-staining TEM. This research used resources of the Advanced Photon Source, a U.S. Department of Energy (DOE) Office of Science User Facility operated for the DOE Office of Science by Argonne National Laboratory under Contract No. DE-AC02-06CH11357. Work at the Molecular Foundry was supported by the Office of Science, Office of Basic Energy Sciences, of the U.S. Department of Energy under Contract No. DE-AC02-05CH11231. This work was supported by the National Institute of Diabetes and Digestive and Kidney Diseases of the NIH under award number R01DK042667 to N.M.D. and supported in part by Institutional Research Grant #16-183-31 from the American Cancer Society and the MCW Cancer Center through a pilot research grant to L.J.O. The content is solely the responsibility of the authors and does not necessarily represents the official views of the NIH.

## Author contributions

N.M.D. and L.J.O. initiated the project. L.J.O. performed mutagenesis, protein expression, purification, crystallization, X-ray data collection (PDB 6P8I) processing (PDB 6P8I), determined structures and structural analysis. K.B. collected and processed the data for the pH 7.0 structure (PDB 6V02). L.J.O conducted SPR experiments and processed/analyzed SPR data, analyzed SAXS data, conducted and analyzed SEC data. G.R. analyzed and supervised collection of negative stain and EM data. S.K.M. collected and analyzed the FPOP data under the supervision of J.S. Modeling done by O.C.G. under supervision of R.J.W. Mass spectrometry done by M.I. supervised by M.T. All authors contributed to data interpretation and preparation of the manuscript. Initial manuscript written by L.J.O. with later versions edited by N.M.D and J-J.P.K. N.D., J-J.P.K and L.J.O. orchestrated the project.

