## Supplemental Table 1 for "Allosteric regulation of lysosomal enzyme recognition by the cation-independent mannose 6-phosphate receptor"

**Extended Data Table 1: Data collection and refinement statistics.**

| | domains1-5 pH 5.5 | domains1-5 pH 7.0 |
|---|---|---|
| Wavelength | 1.54 | 1 |
| Resolution range | 39.97 - 2.54 (2.63 - 2.54) | 35.89 - 2.8 (2.9 - 2.8) |
| Space group | P 1 21 1 | P 1 21 1 |
| Unit cell | 50.788 66.228 123.07<br>90 99.18 90 | 50.78 66.22 124.53<br>90 100.533 90 |
| Total reflections | 99803 (8816) | 36270 (1693) |
| Unique reflections | 26696 (2410) | 18342 (851) |
| Multiplicity | 3.7 (3.4) | 2.0 (2.0) |
| Completeness (%) | 98.37 (91.03) | 90.52 (42.44) |
| Mean I/sigma(I) | 18.41 (2.13) | 14.78 (1.87) |
| Wilson B-factor | 55.14 | 62.20 |
| R-merge | 0.07203 (0.6244) | 0.05846 (0.5394) |
| R-meas | 0.0836 (0.7367) | 0.08267 (0.7629) |
| R-pim | 0.04202 (0.3854) | 0.05846 (0.5394) |
| CC1/2 | 0.997 (0.753) | 0.995 (0.403) |
| CC* | 0.999 (0.927) | 0.999 (0.758) |
| Reflections used in refinement | 26362 (2394) | 18342 (851) |
| Reflections used for R-free | 1285 (114) | 928 (47) |
| R-work | 0.2322 (0.3213) | 0.2402 (0.3890) |
| R-free | 0.2980 (0.3591) | 0.3078 (0.4044) |
| CC(work) | 0.903 (0.646) | 0.892 (0.473) |
| CC(free) | 0.886 (0.639) | 0.796 (0.457) |
| # non-hydrogen atoms | 5306 | 4570 |
| macromolecules | 5217 | 4482 |
| N-linked glycan | 61 | 14 |
| solvent | 28 | 74 |
| Protein residues | 666 | 573 |
| RMS(bonds) | 0.007 | 0.007 |
| RMS(angles) | 1.30 | 1.28 |
| Ramachandran favored (%) | 85.45 | 83.24 |
| Ramachandran allowed (%) | 11.30 | 12.57 |
| Ramachandran outliers (%) | 3.25 | 4.19 |
| Rotamer outliers (%) | 5.91 | 6.41 |
| Clashscore | 6.88 | 8.75 |
| Average B-factor | 37.67 | 45.20 |
| macromolecules | 36.11 | 44.93 |
| carbohydrate | 160.40 | 92.57 |
| solvent | 61.22 | 52.63 |
| PDB ID | 6P8I | 6V02 |

Statistics for the highest-resolution shell are shown in parentheses.
