## extended data legend Fig.1 for "Allosteric regulation of lysosomal enzyme recognition by the cation-independent mannose 6-phosphate receptor"

### Extended Data Fig. 1

**a**

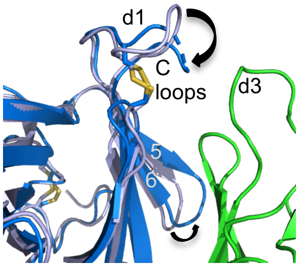


**b**

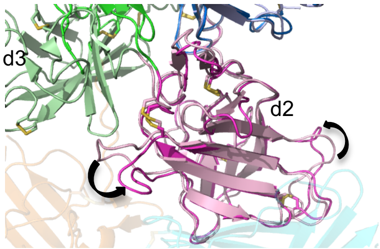


**c**

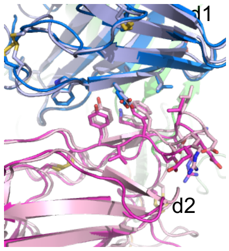


**d**

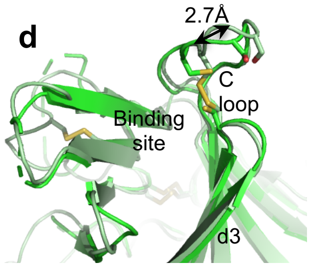


**e**

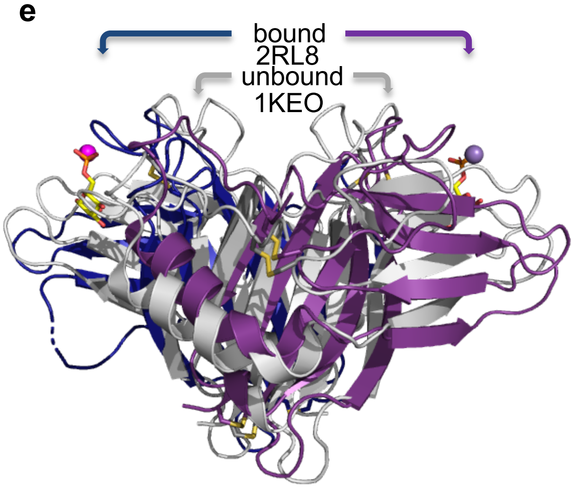


**f**

| Domains | Area (Å²) | CSS* | # H-bonds | #salt bridges |
|---|---|---|---|---|
| 1:3 (unbound)<br>(bound) | 1019<br>370 | 1.0<br>0.30 | 10<br>8 | 2<br>3 |
| 2:5 | 967 | 0.0 | 19 | 10 |
| 1:2 (unbound)<br>(bound) | 782<br>860 | 1.0<br>1.0 | 8<br>8 | 3<br>2 |
| 2:4 | 503 | 0.0 | 4 | 1 |
| 4:5 | 477 | 0.0 | 6 | 6 |
| 2:3 (unbound)<br>(bound) | 262<br>717 | 0.097<br>0.79 | 8<br>18 | 4<br>4 |
| CD-MPR(1KEO)<br>2RL8 | 769<br>1044 | 1.0<br>0.46 | 11<br>9 | 0<br>2 |

g

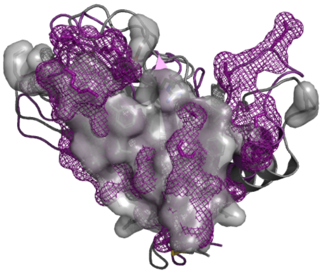

**h**

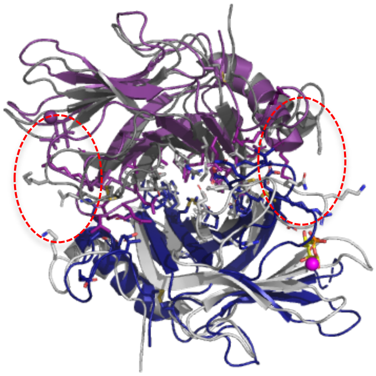
