## Supplemental Fig. 1 for "Allosteric regulation of lysosomal enzyme recognition by the cation-independent mannose 6-phosphate receptor"

**Extended Data Fig. 1. Influence of ligand presence on members of the P-type lectin family. a,** Movement of CI-MPR domain 1 loop C in the presence (light blue) and absence of ligand (blue). **b,** Loop movements in CI-MPR domain 2 in the presence (light pink) and absence of ligand (magenta). **c,** Shifting of amino acid positions at the interface of CI-MPR domains 1 and 2 in the presence (light pink) and absence of ligand (magenta)**. d,** Loop C of CI-MPR domain 3 alters position depending on the presence (light green) or absence of ligand (green). **e,** CD-MPR in bound (dark purple) and unbound (grey) forms. **f,** Table of interface areas and CSS values calculated with PISA. Comparison of interface of CD-MPR in bound (purple) and unbound (grey) highlighting differences in areas **(g)** and salt bridges/H-bonding **(h)**.
