## Supplementary figures and images for "Allosteric regulation of lysosomal enzyme recognition by the cation-independent mannose 6-phosphate receptor"

### extended data legend Fig.2

#### Extended Data Fig. 2

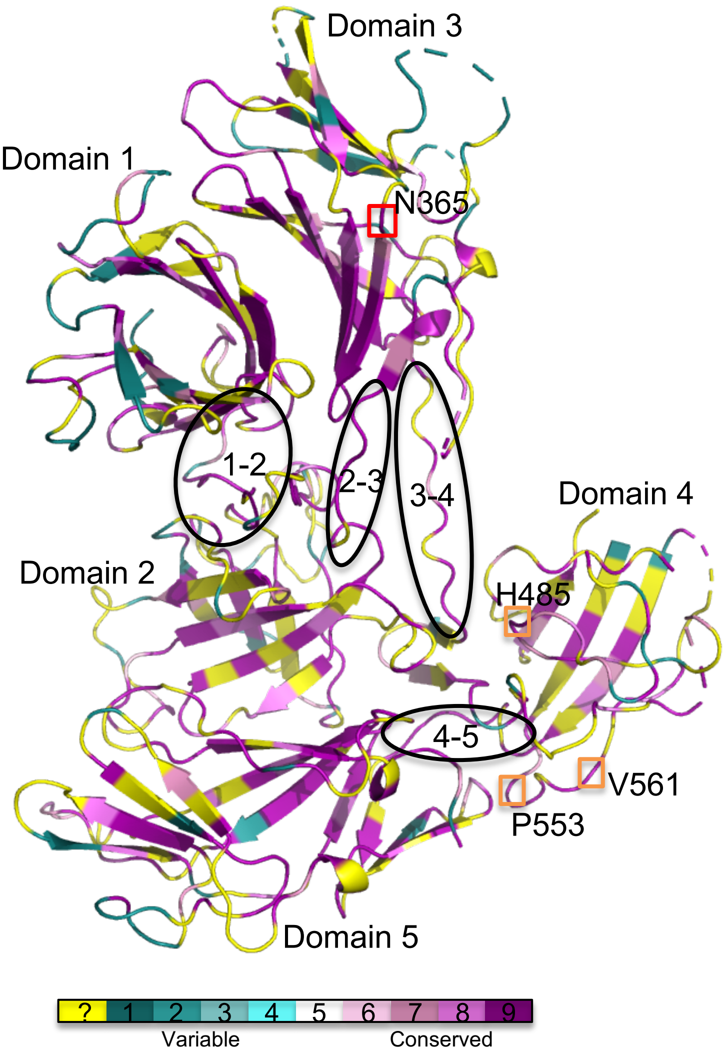

### extended data legend Fig.3

#### Extended Data Fig. 3

**a**

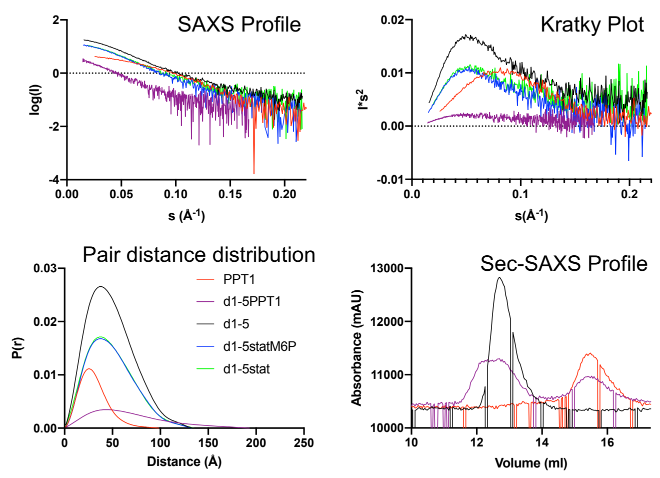


**b**

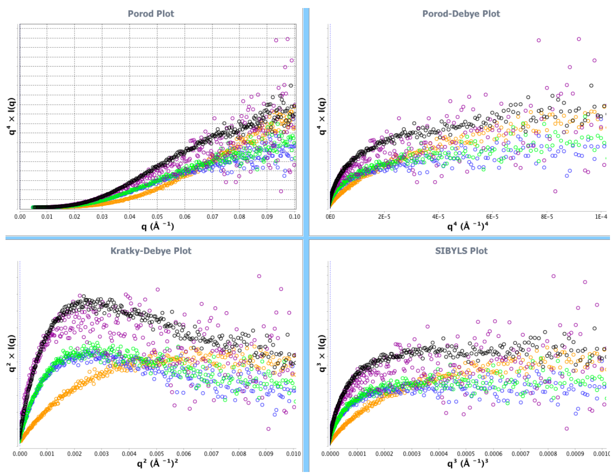


**c**

| | $R_g$ | $D_{max}$ | $P_E$ | Vol | MW | AA MW* |
|---|---|---|---|---|---|---|
| | (Å) | (Å) | | (Å³) | kDA | |
| D1-5stat | 37 | 101 | 2.9 | 140000 | 87.5 | 81.3 |
| D1-5statM6P | 37 | 102 | 2.9 | 156000 | 97.5 | 81.3 |
| D1-5 | 38 | 115 | 2.9 | 145000 | 90.6 | 81.3 |
| PPT1 | 23 | 59 | 1.3 | 59900 | 37.4 | 31.3 |
| D1-5+PPT1 | 41 | 114 | 2.8 | 203000 | 126.9 | 112.6 |

**d**

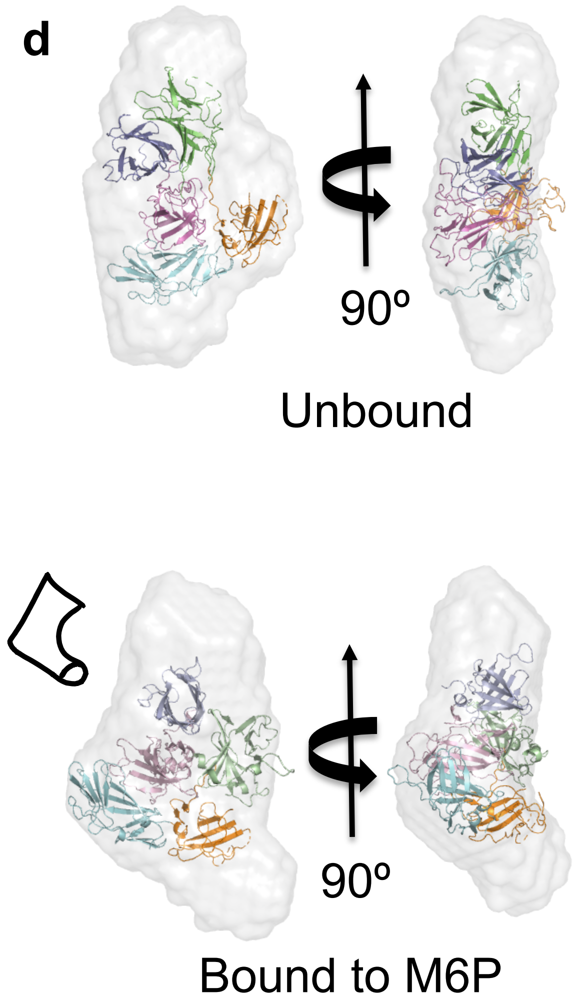


**e**

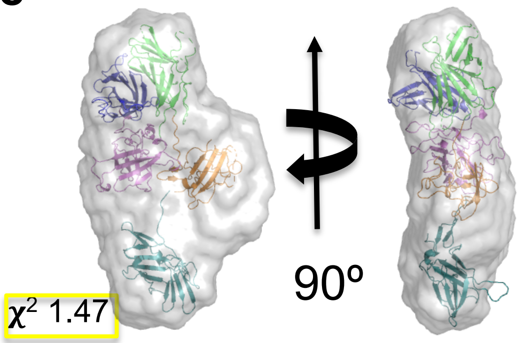

### extended data legend Fig.4

#### Extended Data Fig. 4

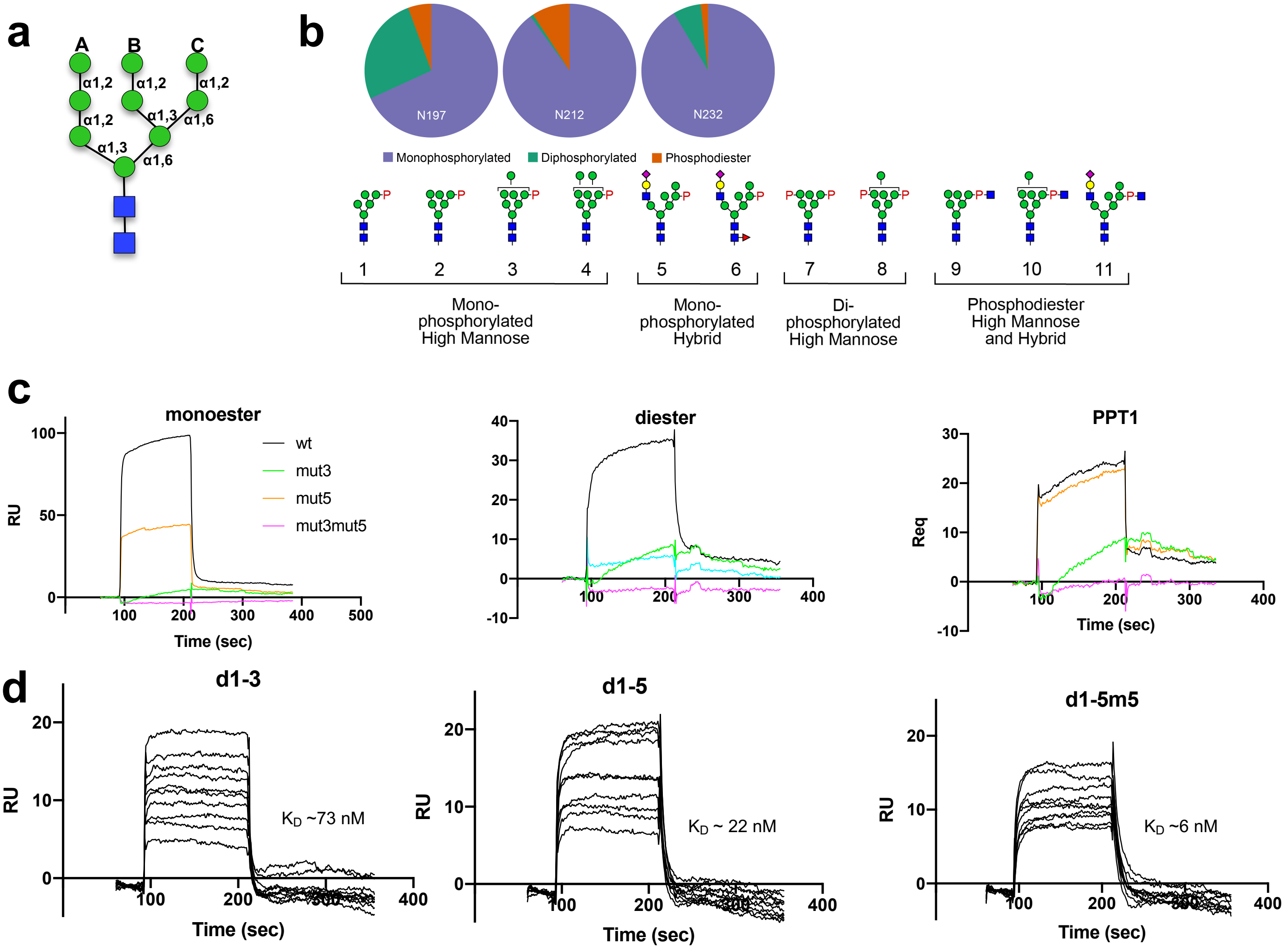

### extended data legend Fig.5

#### Extended Data Fig. 5

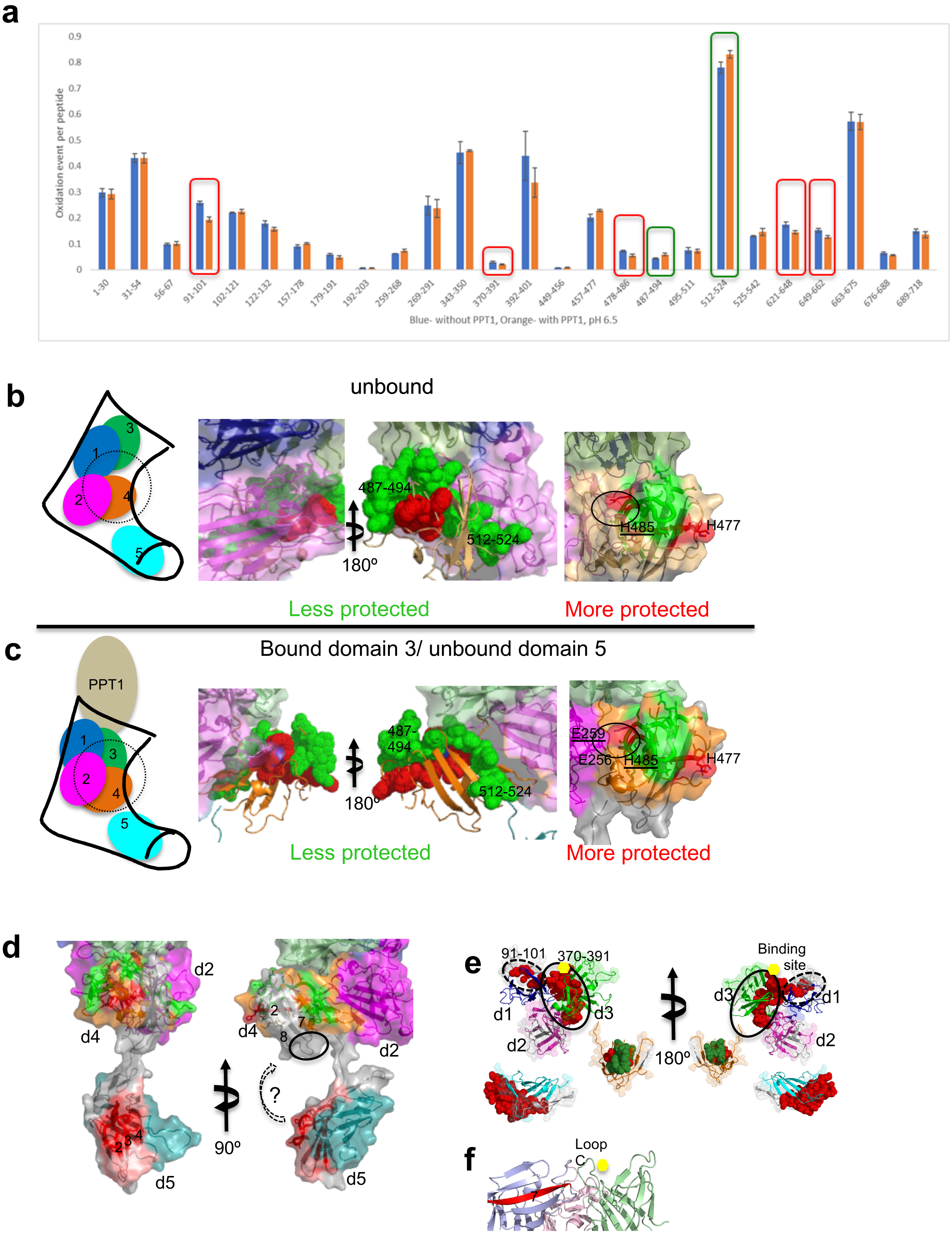

### extended data legend Fig.6

#### Extended Fig. 6

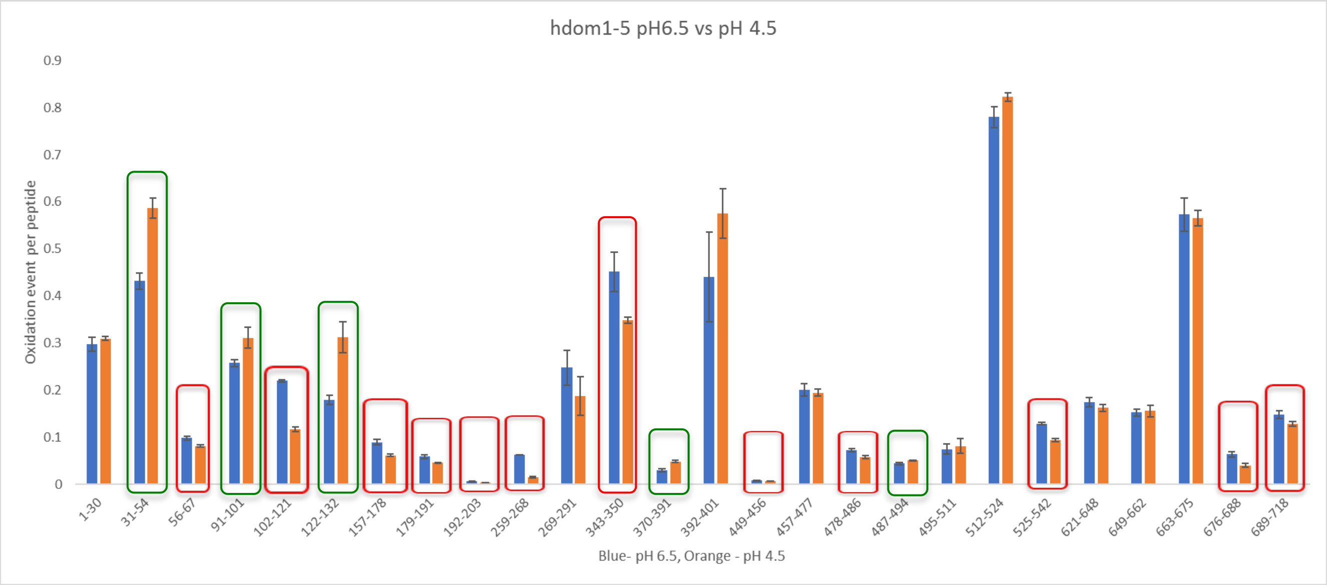

### extended data legend Fig.7

#### Extended Fig. 7

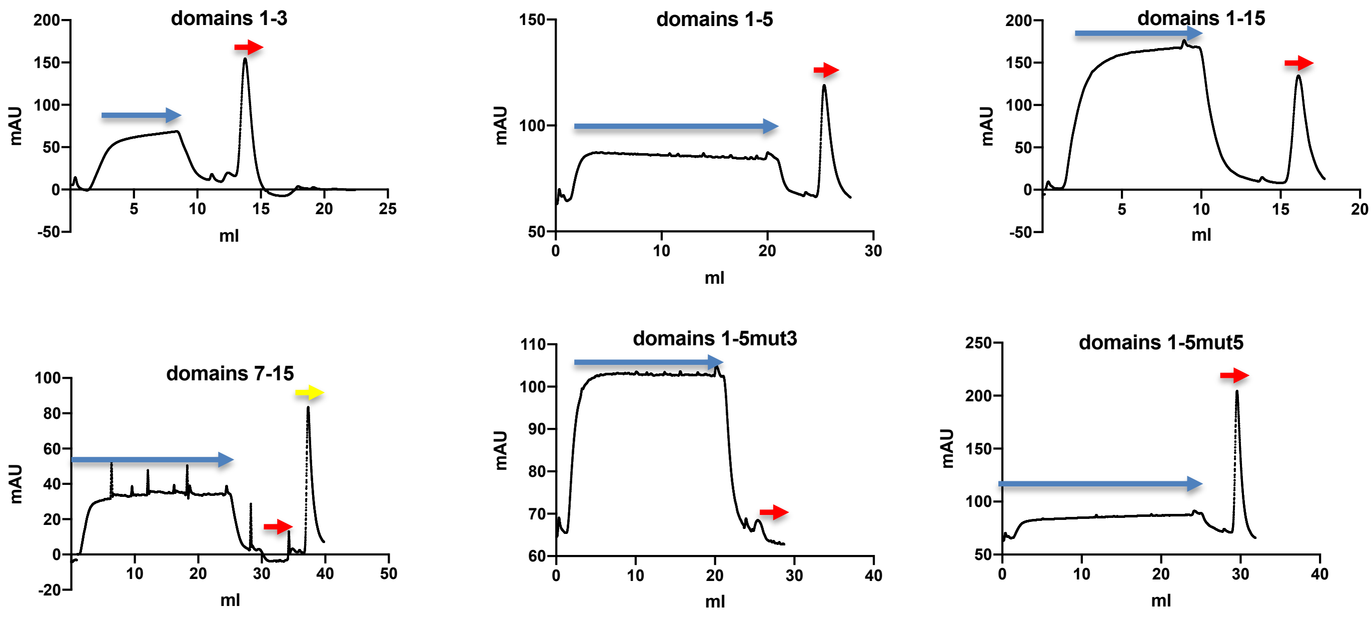
