## Supplemental Fig 2 for "Allosteric regulation of lysosomal enzyme recognition by the cation-independent mannose 6-phosphate receptor"

**Extended Data Fig. 2. Consurf results showing amino acid species conservation.** The PDB 6P8I structure was analyzed via the ConSurf server to determine regions of amino acid conservation and then mapped onto the model^1^. The individual domains of domains 1-5 are labeled while the linkers are labeled and circled. Amino acids are colored by degree of conservation as shown (conserved, purple; variable, cyan).
