## Supplemental Fig 3 for "Allosteric regulation of lysosomal enzyme recognition by the cation-independent mannose 6-phosphate receptor"

**Extended Data Fig. 3. SAXS data plots. a,** Plots of the SAXS Profile, Kratky Plot, Pairwise distance distribution (SEC-SAXS: domains 1-5 black, PPT1 orange, and complex of CI-MPR domains 1-5 with PPT1 purple; batch mode samples: domains 1-5 green, domains 1-5 with M6P, blue). Lower right panel of (a) is elution profile of domains 1-5 with PPT1 from a G200-Increase column. **b**, Flexibility plots colored as in panel (a) above. **c**, table of SAXS derived values. **d**, cuvette collected data for, upper, PDB 6P8I crystal structure modeled into SAXS derived envelope for domains 1-5 and lower, modified domain 3 structure inserted into envelope derived from data collected on domains 1-5 in the presence of M6P.
