## Supplemental Fig 4 for "Allosteric regulation of lysosomal enzyme recognition by the cation-independent mannose 6-phosphate receptor"

**Extended Data Fig. 4. Characterization of glycans of recombinant PPT1 by Mass spectrometry. a,** Schematic illustration of a typical high mannose glycan with linkages and arms are labeled. Green circles represent mannose residues while blue square represent N-acetylglucosamine residues. **b**, Pie charts depicting percentages of glycans found in recombinant PPT1 protein above schematic diagram representing various forms of glycans identified. **c**, SPR sensorgrams for either domains 1-5 truncated protein or CRD binding mutants (mut3, domains 1-5R391A^2^; mut5, domains 1-5RR688A^3^; mut3mut5, domains1-5R391A/R688A) (120 nM) flowing over GAA-phosphomonoester **(upper left)** or GAA-monodiester **(upper center)** and PPT1 **(upper right)** surfaces. Mutation of domain 5 has little to no effect on domains 1-5 ability to bind PPT1 but eliminates the protein’s ability to bind to the diester surface. **d**, Sensorgrams for domains 1-3, domains 1-5 or domains 1-5mut5 truncated protein (10 to 120 nM) flowing over PPT1 surface.
