## Supplemental Fig 5 for "Allosteric regulation of lysosomal enzyme recognition by the cation-independent mannose 6-phosphate receptor"

**Extended Data Fig. 5. The FPOP results (a, tryptic peptide fragments identified in the absence (blue) and presence (orange) of PPT1) were mapped onto our SAXS models representing domains 1-5 in the unbound and bound domain 3 conformation.** Peptides undergoing statistically significant (p≤0.05) changes between the two conditions are boxed (red, more protected in presence PPT1; green, less protected in the presence of PPT1). **b,** Looking first at the changes in protection for domain 4 we can see that these models account for the deprotection of peptides 487-494 and 512-524 upon ligand binding: going from a protected interface with domain 3 in the unbound position to a more solvent exposed position in the bound structure. Although peptide 478-488 on domain 4 could be a candidate to interact with domain 5, close inspection indicates this is unlikely. Only N-terminal residues H478, F479 and F480 and C-terminal residues H485 and R486 are solvent exposed in the unbound model with the three N-terminal residues remaining solvent exposed in the bound model **(c)**. However, species conserved H485 becomes part of the interface with domain 2 forming a hydrogen bond with mainly conserved E256 and is most probably accounts for the change in reactivity of this peptide **(c, right panel)**. **d**, Domain 5 has two peptides which show protection from oxidation in the presence of ligand compared to absence (peptides 621-648 (strands 2 and 3) and 649 to 662 (strand 4)) while domain 4 has one peptide (478-488, stand 2). However, species conserved H485 becomes part of the interface with domain 2 forming a hydrogen bond with mainly conserved E256 and is most probably accounts for the change in reactivity of this peptide **(c, right panel)**. FPOP did not provide data on all regions of the domain 1-5 protein and in particular areas of domain 4 and domain 5 which could be in position to interact upon ligand binding: C-terminal β-sheet of domain 4. A likely candidate for interaction with domain 5 is the loop between strands 7 and 8 of domain 4 (P553-V561). It is highly species conserved and is not currently known to be involved in any domain-domain contacts but is poised to interact with domain 5 (621-648, 649-65) **(d)**. The two other peptides with altered protection in the presence of ligand are located in domains 1 and 3 **(e)**. Peptide 370-391 is located in the binding site of domain 3 and encompasses the previously identified highly conserved, essential for ligand binding residue R391^4^. The peptide 91-101 is located on strand 7 on the back sheet of domain 1 and when domain 3 is in the bound position, this strand is across from the ligand binding site (yellow hexagon) **(f)**. Although this peptide is outside the known ligand binding region and has not previously been associated with any functions, its proximity to the known binding site makes it worthwhile investigating as a secondary site of ligand interactions.
