## Supplemental Fig 6 for "Allosteric regulation of lysosomal enzyme recognition by the cation-independent mannose 6-phosphate receptor"

**Extended Data Fig. 6. The FPOP results (a, tryptic peptide fragments identified at pH 6.5 (blue) and pH 4.5 (orange).** Peptides undergoing statistically significant (p≤0.05) changes between the two conditions are boxed (red, more protected at pH 4.5; green, less protected at pH 4.5).
