## Supplemental Fig 7 for "Allosteric regulation of lysosomal enzyme recognition by the cation-independent mannose 6-phosphate receptor"

**Extended Data Fig. 7. Chromatograms of CI-MPR constructs on PPT1 affinity column.** Blue arrows indicate loading of CI-MPR sample onto 1 ml PPT1 affinity column generated by primary amine coupling. Red arrow indicates elution of column with pH 4.5 buffer and yellow arrow indicates elution of column with buffer containing 10 mM M6P.
